# Tsc1-mTOR signaling controls the structure and function of midbrain dopamine neurons

**DOI:** 10.1101/376814

**Authors:** Polina Kosillo, Natalie M. Doig, Alexander H.C.W. Agopyan-Miu, Kamran Ahmed, Lisa Conyers, Sarah Threlfell, Peter J. Magill, Helen S. Bateup

**Affiliations:** Department of Molecular and Cell Biology, University of California, Berkeley, Berkeley, CA, 94720, USA; Medical Research Council Brain Network Dynamics Unit, University of Oxford, Oxford OX1 3TH, United Kingdom; Department of Physiology, Anatomy and Genetics, University of Oxford, Oxford OX1 3PT, United Kingdom; Oxford Parkinson’s Disease Centre, University of Oxford, Oxford, OX1 3QX, United Kingdom; Helen Wills Neuroscience Institute, University of California, Berkeley, Berkeley, CA 94720, USA; Lead contact

**Keywords:** mTOR complex 1, Tsc1, Raptor, dopamine neurons, dopamine release, voltammetry, electron microscopy, cognitive flexibility, Tuberous Sclerosis Complex, autism spectrum disorder

## Abstract

mTOR complex 1 (mTORC1) is a central coordinator of cell growth and metabolism. Mutations in regulators of mTORC1 cause syndromic disorders with a high prevalence of cognitive and psychiatric conditions. To elucidate the cellular origins of these manifestations, we conditionally deleted the gene encoding the mTORC1 negative regulator Tsc1 from mouse midbrain dopamine neurons, which modulate motor, affective, and cognitive behaviors that are frequently affected in psychiatric disorders. Loss of Tsc1 and constitutive activation of mTORC1 strongly impacted the properties of dopamine neurons, causing somatodendritic hypertrophy, reduced intrinsic excitability, altered axon terminal ultrastructure, and severely impaired dopamine release. These perturbations were associated with selective deficits in cognitive flexibility, which could be prevented by genetic reduction of the obligatory mTORC1 protein Raptor. Our results establish a critical role for mTORC1 in setting the functional properties of midbrain dopamine neurons, and indicate that dopaminergic dysfunction may underlie cognitive inflexibility in mTOR-related syndromes.

## Introduction

The mechanistic target of rapamycin (mTOR) pathway is a highly conserved, fundamental signaling cascade that integrates intra- and extracellular signals to regulate a variety of cellular metabolic processes. Deregulation of mTOR signaling is linked to numerous diseases, with particularly detrimental outcomes for nervous system development and function (Costa-Mattioli and Monteggia, 2013; Lipton and Sahin, 2014). Precisely how perturbed mTOR signaling affects the activity of neurons and neural circuits is a key open question. mTOR functions as a kinase within two complexes, mTORC1 and mTORC2, that have distinct protein components, upstream activators, and downstream targets (Hoeffer and Klann, 2010). Activation of mTORC1 via PI3K/Akt signaling promotes anabolic functions including protein and lipid synthesis, and suppresses catabolic processes such as autophagy, leading to cell growth and division (Huang and Manning, 2008). mTORC1 activity is negatively regulated by the heterodimeric Tsc1/2 protein complex that exerts GTP-ase activating (GAP) activity on the small GTP-ase Rheb, a direct activator of mTORC1 (Laplante and Sabatini, 2012). Disruption of the Tsc1/2 complex results in constitutively active mTORC1 leading to altered cell growth, metabolism, and proliferation (Huang and Manning, 2008).

In post-mitotic neurons, Tsc1/2-mTORC1 signaling controls neuronal communication via regulation of membrane excitability, synaptic transmission, and synaptic plasticity (Bateup et al., 2011, 2013; Benthall et al., 2018; Ehninger et al., 2008; Normand et al., 2013; Tavazoie et al., 2005; Tsai et al., 2012; Weston et al., 2014; Yang et al., 2012). Loss-of-function mutations in negative regulators of mTORC1 cause neurodevelopmental syndromes associated with benign tumors in multiple organs and significant neurological and psychiatric impairments. In particular, patients with Tuberous Sclerosis Complex (TSC), caused by mutations in *TSC1* or *TSC2*, frequently present with neuropsychiatric conditions including autism spectrum disorder (ASD), attention deficit hyperactivity disorder (ADHD), anxiety, and aggression, which are collectively termed TAND (TSC-Associated-Neuropsychiatric-Disorders) (de Vries et al., 2015). In contrast to the well-studied mechanisms of tumor formation in TSC, less is known about how mutations in *TSC1/2* and deregulation of mTORC1 signaling cause neuropsychiatric and behavioral abnormalities. In particular, the specific neuronal populations responsible for TAND and the functional consequences of mTOR-pathway mutations on these neurons remain to be defined.

Midbrain dopamine (DA) neurons of the substantia nigra pars compacta (SNc) and ventral tegmental area (VTA) modulate a host of behaviors and functions including movement, cognition, and reward learning (Schultz, 2005; Steinberg et al., 2013; Wickens et al., 2007). Given the involvement of DA signaling in many of the psychiatric conditions associated with TSC and other mTOR-related disorders, we hypothesized that changes in DA signaling may be central to TAND. To test this, we selectively deleted *Tsc1* from mouse DA neurons and investigated how developmental deregulation of mTORC1 affects midbrain DA neuron function. This approach also allowed us to test whether activation of mTORC1 signaling in DA neurons alone was sufficient to drive TAND-related behavioral phenotypes.

We found that deletion of *Tsc1* from DA neurons profoundly affected their structural and functional properties resulting in significant impairments in DA release. This dopaminergic deficit led to reduced cognitive flexibility in the absence of changes to motor, social, or affective behaviors. Our findings identify mTORC1 as a critical regulator of midbrain DA neuron output and suggest that cognitive flexibility deficits in mTOR-related disorders may be driven by changes in DA signaling.

## Results

### Loss of Tsc1 from DA neurons causes somatic and dendritic hypertrophy

To selectively activate mTORC1 signaling in DA neurons, we conditionally deleted *Tsc1*, which results in loss of function of the Tsc1/2 complex (Kwiatkowski et al., 2002) (Figure S1A), from DA neurons using *DAT^IRES^Cre* mice (Bäckman et al., 2006). For all experiments, mice were heterozygous for *DAT^IRES^Cre* and had either wild-type (“DA-Tsc1 WT”) or homozygous floxed alleles of *Tsc1* (“DA-Tsc1 KO”) (Figure S1B). To confirm that loss of Tsc1 resulted in functional activation of mTORC1 signaling in midbrain DA neurons, we quantified phosphorylation of the mTORC1 pathway target ribosomal protein S6 (p-S6) in tyrosine hydroxylase (TH)-labeled SNc and VTA neurons (Figures 1A-D and S1C). p-S6 levels were significantly higher in DA-Tsc1 KO neurons compared to DA-Tsc1 WT (Figures 1E and S1D), indicative of activated mTORC1 signaling. Consistent with the known function of mTORC1 in controlling cell size (Lipton and Sahin, 2014), loss of Tsc1 caused a profound increase in DA neuron soma area (Figures 1F and S1E).

**Figure 1.**
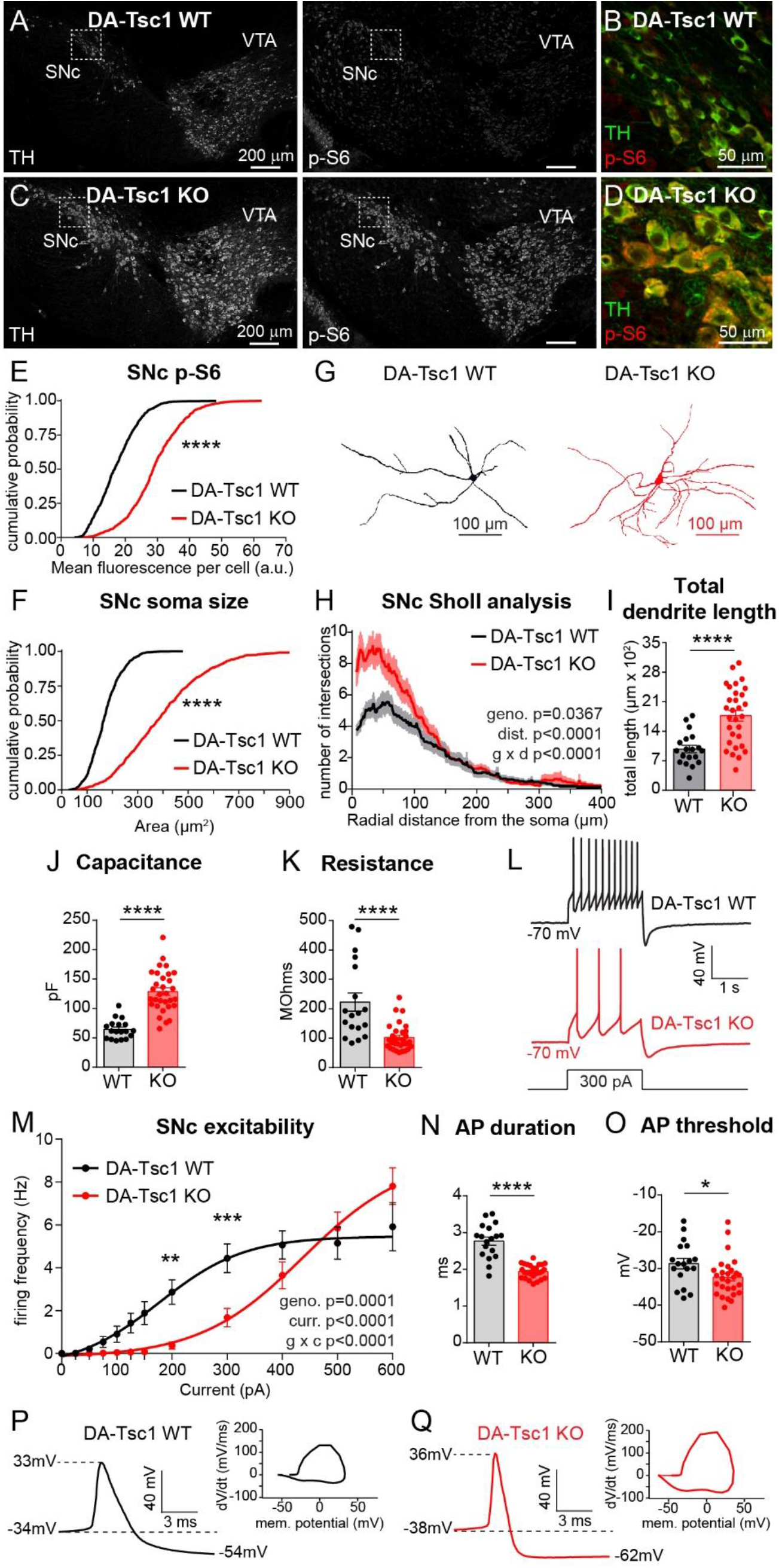
DA-Tsc1 KO SNc neurons are hypertrophic and have reduced intrinsic excitability. (A-D) Confocal images of coronal midbrain sections from *DAV^IRES^Cre^wt/+^* mice homozygous for wild-type (DA-Tsc1 WT, A-B) or floxed *Tsc1* (DA-Tsc1 KO, C-D). Sections were labeled with antibodies against tyrosine hydroxylase (TH) and phosphorylated S6 (p-S6, Ser240/244). Panels B and D show higher magnification merged images of the boxed regions in A and C. SNc=substantia nigra pars compacta, VTA=ventral tegmental area. (E,F) Cumulative probability plots of SNc DA neuron p-S6 levels (E) and soma area (F). Black lines show the distributions of values from DA-Tsc1 WT mice (n=759 neurons from 3 mice) and red lines show distributions from DA-Tsc1 KO mice (n=855 neurons from 3 mice). ****, p<0.0001, Kolmogorov-Smirnov tests. (G) Three-dimensional reconstructions of the soma and dendrites of neurobiotin-filled SNc DA neurons from whole-cell patch-clamp experiments. (H) Sholl analysis of SNc DA neurons. Dark colored lines are the mean, the lighter color shading is SEM (DA-Tsc1 WT: n=19 neurons from 5 mice, DA-Tsc1 KO: n=30 neurons from 9 mice). Two-way ANOVA p values are shown. (I) Total dendritic length per cell measured from reconstructed SNc DA neurons. Bars represent mean ± SEM, dots represent individual neurons. (n is the same as for panel H) ****, p<0.0001, unpaired, two-tailed t test. (J,K) Bar graphs display mean ± SEM membrane capacitance (J), and membrane resistance (K). Dots indicate the values of individual neurons. (DA-Tsc1 WT: n=18 neurons from 5 mice, DA-Tsc1 KO: n=27 neurons from 9 mice). ****, p<0.0001, unpaired, two-tailed t tests. (L) Typical examples of action potential firing elicited with a 300 pA current step in SNc DA neurons of the indicated genotypes. (M) Excitability curves showing the firing frequency of SNc DA neurons in response to two second depolarizing current steps of increasing amplitude. Data are displayed as mean ± SEM (DA-Tsc1 WT: n=20 neurons from 5 mice, DA-Tsc1 KO: n=32 neurons from 9 mice). Two-way ANOVA p values are shown. **, p=0.0014; ***, p=0.0002, Sidak’s multiple comparisons test. (N,O) Bar graphs display mean ± SEM action potential (AP) duration (N), and membrane potential at AP threshold (O). Dots indicate the values of individual neurons. (DA-Tsc1 WT: n=18 neurons from 5 mice, DA-Tsc1 KO: n=27 neurons from 9 mice). *, p=0.0338; ****, p<0.0001, unpaired, two-tailed t tests. (P,Q) Examples of individual action potentials evoked by positive current injection and their respective phase plots for SNc DA neurons in DA-Tsc1 WT (P) and DA-Tsc1 KO mice (Q). See also Figure S1 and Tables S1 and S2 for complete electrophysiology results.

In addition to somatic hypertrophy, altered dendrite branching has been observed in neurons with mutations in mTORC1 regulators (Benthall et al., 2018; Goto et al., 2011; Kwon et al., 2006; Weston et al., 2014). To investigate how loss of Tsc1 affects the dendrites of DA neurons, we performed Sholl analysis on three-dimensional reconstructions of neurobiotin-filled neurons (Figure S1I). We found that SNc and VTA DA neurons in DA-Tsc1 KO mice had more complex dendrites and increased total dendritic length (Figure 1G-I and S1F-H). Compared to WT neurons, the dendrites of SNc DA-Tsc1 KO neurons extended about the same radial distance from the soma (Figure 1H), but were longer in VTA DA-Tsc1 KO neurons (Figure S1G). Together these data demonstrate that loss of Tsc1, and constitutive activation of mTORC1 signaling, strongly impact the somatodendritic architecture of midbrain DA neurons.

### DA neurons exhibit reduced intrinsic excitability and altered action potential shape following deletion of *Tsc1*

The structural properties of neurons dictate their function. Therefore, we tested whether the extensive somatodendritic remodeling we observed in DA-Tsc1 KO neurons resulted in a change to their intrinsic excitability. To visualize DA neurons in acute slices for electrophysiology recordings, we bred DA-Tsc1 KO mice to ‘reporter mice’ expressing Cre-dependent tdTomato (Ai9 line; Madisen et al., 2010) (Figure S1B). Consistent with their somatodendritic hypertrophy, SNc and VTA DA-Tsc1 KO neurons had significantly increased membrane capacitance and decreased membrane resistance compared to WT neurons (Figures 1J,K and S1J,K and Tables S1 and S2). We injected steps of depolarizing current to generate excitability curves and found that, consistent with the changes in passive membrane properties, SNc and VTA DA-Tsc1 KO cells were hypo-excitable compared to DA-Tsc1 WT (Figures 1L,M and S1 L,M and Tables S1 and S2). In particular, significantly larger current step amplitudes were required to evoke action potential (AP) firing in DA-Tsc1 KO neurons (Figure 1M and S1M). At current amplitudes up to 400 pA, DA-Tsc1 KO neurons fired fewer APs than DA-Tsc1 WT neurons; however, at the 500 and 600 pA current steps, DA-Tsc1 KO neurons had similar firing rates as DA-Tsc1 WT cells (Figures 1M and S1M). This suggests that higher currents are required to drive increased spiking in DA-Tsc1 KO neurons but that these neurons may still be able to fire a ‘burst’ of APs given a sufficiently large excitatory input.

DA neurons have been classically defined by their relatively long AP duration and prominent after-hyperpolarization (Bean, 2007). A variety of sodium, calcium and potassium channels control DA neuron firing properties (Gantz et al., 2018). Since mTORC1 signaling can regulate ion channel expression (Raab-Graham et al., 2006; Yang et al., 2012), we examined the properties of single APs evoked by positive current injection in DA-Tsc1 KO neurons. *Tsc1* deletion caused a substantial change in the AP waveform of DA neurons (Figures 1N-Q and S1N-Q). Specifically, the AP duration of SNc and VTA DA-Tsc1 KO neurons was significantly shorter and threshold was reduced in SNc neurons compared to WT (Figures 1N,O and S1N,O and Tables S1 and S2). These changes are indicative of altered potassium channel conductances that control repolarization (Kimm et al., 2015). Notably, these results are similar to those of a prior study that reported shorter duration APs in *Tsc1* knock-out thalamic neurons (Normand et al., 2013).

### Loss of Tsc1 causes severe impairments in striatal DA release

The AP firing of DA neurons does not linearly translate into axonal DA release events (Rice et al., 2011; Sulzer et al., 2016). Therefore, a direct interrogation of DA release is necessary to determine how mTORC1 hyperactivity impacts dopaminergic neurotransmission. To this end, we used fast-scan cyclic voltammetry (FCV) at carbon fiber microelectrodes (CFM) to monitor electrically-evoked DA release in the striatum, the brain region with the most dense dopaminergic innervation. To directly compare DA transients between the two genotypes, we performed experiments with paired striatal slices from DA-Tsc1 WT and DA-Tsc1 KO mice with interleaved recordings using the same CFM.

Sampling of extracellular dopamine release ([DA]_o_) evoked by electrical stimulation at different striatal locations revealed significant reductions in peak [DA]_o_ amplitudes throughout the dorsal striatum, but not in the nucleus accumbens core (p = 0.1650, paired t-test), in DA-Tsc1 KO mice compared to DA-Tsc1 WT (Figures 2A,B). Decreased DA release was found with both single pulse (Figure 2A,B) and high-frequency burst stimulation (4 pulses at 100 Hz, Figure S2A,B). Because peak-evoked [DA]_o_ in the nucleus accumbens was not significantly altered by *Tsc1* deletion, we focused subsequent analyses on the dorsal striatum. Combining all dorsal striatal sites per mouse, we found an average 60% reduction in single pulse (Figure 2C,D) and 45% reduction in high-frequency burst-evoked [DA]_o_ (Figure S2C,D) following *Tsc1* deletion. We tested multiple stimulation intensities and found that DA release deficits in DA-Tsc1 KO slices were exacerbated at low stimulation intensities, consistent with the intrinsic hypoexcitability observed at the cell body, while peak-evoked [DA]_o_ plateaued at similar stimulation intensities as in WT (Figure S2E).

**Figure 2.**
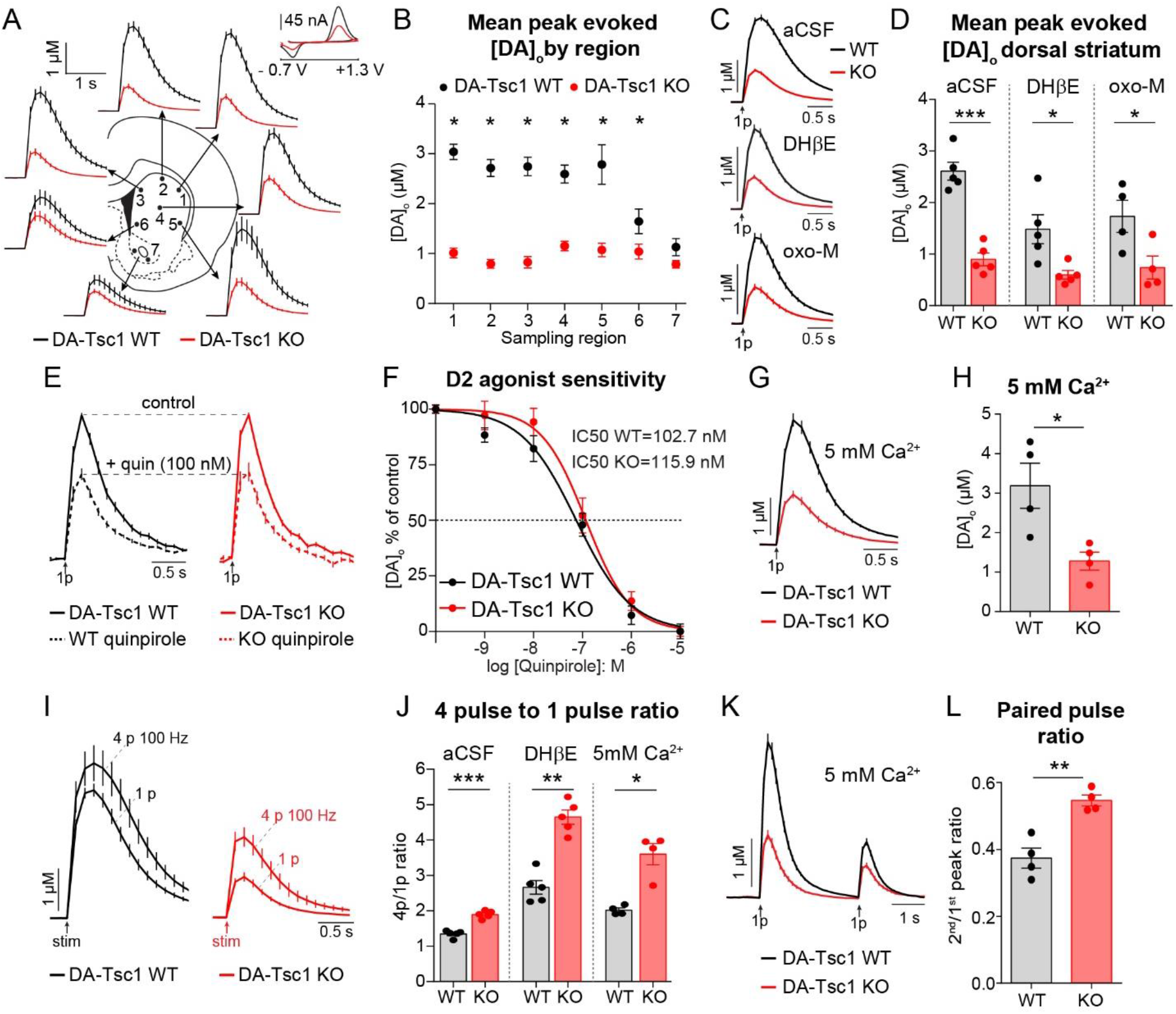
Loss of Tsc1 causes impairments in striatal DA release. (A) Mean extracellular DA release ([DA]_o_) ± SEM versus time evoked from different striatal subregions by single electrical stimuli. Traces are an average of 16-20 release transients per site from 5 mice per genotype. 1-dorsolateral striatum, 2-dorsocentral striatum, 3-dorsomedial striatum, 4-central striatum, 5-ventrolateral striatum, 6-ventromedial striatum, 7-nucleus accumbens core (two sampling sites within the core were averaged together). Inset, typical cyclic voltammograms show characteristic DA waveform. (B) Mean peak single pulse-evoked [DA]_o_ ± SEM by striatal region (numbers correspond to the numbered sites in panel A). n=16-20 transients per site from 5 mice per genotype. *, p_1-4_<0.0001; *, p_5_=0.0002; *, p_6_=0.0378, paired t tests. (C) Mean single pulse-evoked [DA]_o_ ± SEM versus time across all dorsal striatum sites recorded in normal aCSF (average of 96 transients across 6 recording sites per genotype from 5 mice), DHβE (1 μM, average of 116 transients across 6 recording sites per genotype from 5 mice), or oxotremorine-M (oxo-M, 10 μM, average of 86 transients across 6 recording sites per genotype from 4 mice). (D) Mean peak single pulse-evoked [DA]_o_ ± SEM averaged across all dorsal striatum sites (sites #1-6 in panels A and B). Dots represent average evoked [DA]_o_ per mouse in aCSF, DHβE (1 μM) and oxotremorine-M (10 μM), n=4-5 mice per genotype. *, p_DHβE_ =0.0126; *, p_oxo-M_=0.0479; ***, p_acsf_=0.0002, paired t tests. (E) Mean single pulse-evoked [DA]_o_ ± SEM versus time in dorsolateral striatum in control conditions (solid lines) and in the presence of the D2 receptor agonist quinpirole (100 nM, dashed lines). Traces are peak normalized to control values prior to drug application within each genotype. All recordings are in the presence of DHβE (1 μM). Average of 18 transients per drug concentration from 3 mice. (F) Mean ± SEM quinpirole dose-response curves for single pulse-evoked [DA]_o_ in dorsolateral striatum, sigmoidal curve-fit. All recordings in the presence of DHβE (1 μM). n=3 mice per genotype. (G) Mean single pulse-evoked [DA]_o_ ± SEM versus time across all dorsal striatum sites in 5 mM extracellular calcium and 1 μM DHβE. Traces are an average of 96 transients across 6 recording sites per genotype from 4 mice. (H) Mean peak single pulse-evoked [DA]_o_ ± SEM in 5 mM extracellular calcium averaged across all dorsal striatum sites. All recordings in the presence of DHβE (1 μM). Dots represent average evoked [DA]_o_ per mouse, n=4 mice per genotype. *, p=0.0128, paired t test. (I) Mean [DA]_o_ ± SEM versus time evoked by single pulse (“1p”) or high-frequency stimulation (4 pulses at 100 Hz, “4p”) in dorsolateral striatum in normal aCSF. Traces are an average of 18 transients for 1p and 9 transients for 4p per genotype from 5 mice. (J) The ratio of peak [DA]_o_ evoked by a 4 pulse train to single pulse stimulation in dorsolateral striatum in aCSF, DHβE (1 μM), or high extracellular Ca^2+^ (5 mM, 1 μM DHβE applied throughout). Dots represent average 4p/1p ratio per mouse, error bars are SEM. n=4-5 mice per genotype. *, p=0.0204; **, p=0.0024; ***, p=0.0009, unpaired t tests. (K) Mean dorsal striatal [DA]_o_ ± SEM versus time evoked by single pulse stimulations (1p) delivered 3 s apart in high extracellular Ca^2+^ (5 mM, 1 μM DHβE applied throughout). Traces are an average of 96 transients across 6 recording sites per genotype from 4 mice. (L) The ratio of peak [DA]_o_ evoked by the second versus the first pulse in high extracellular Ca^2+^ (5 mM, 1 μM DHβE applied throughout). Dots represent average 2^nd^/1^st^ peak ratio per mouse, error bars are SEM. n=4 mice per genotype. **, p=0.0055, paired t test. See also Figure S2.

Striatal DA release is tightly regulated by striatal cholinergic interneurons (ChIs), which control release probability and can drive axonal DA release directly (Rice et al., 2011; Sulzer et al., 2016). Therefore, we repeated the FCV experiments with cholinergic transmission blockers to test whether the DA release deficits in DA-Tsc1 KO mice were cell autonomous. We used dihydro-ß-erythroidine (DHßE) to block ß2-contaning nicotinic acetylcholine receptors (nAChRs) on DA axon terminals or oxotremorine-M (oxo-M) to activate muscarinic autoreceptors on ChIs, which inhibit acetylcholine release (Figure S2F). We found that the single pulse-evoked release deficits in DA-Tsc1 KO mice persisted after blockade of cholinergic transmission (Figure 2C,D). The reduction in high-frequency burst-evoked release in DA-Tsc1 KO mice was more variable in cholinergic blockers (Figure S2C,D). However, this is expected for pulse train stimulation paradigms in which DA release is subject to moment-by-moment variations in release probability due to short-term plasticity (Cragg, 2003; Cragg and Rice, 2004; Montague et al., 2004). Overall, we conclude that cell-autonomous changes are responsible for the profound reduction in evoked [DA]_o_ in the dorsal striatum of DA-Tsc1 KO mice.

### Changes in D2 receptor occupancy or calcium sensitivity do not account for DA release deficits in DA-Tsc1 KO mice

One possible mechanism for a cell autonomous reduction in striatal [DA]_o_ is increased activity or expression of type 2 dopamine (D2) autoreceptors located on DA axons, which inhibit DA release (Sulzer et al., 2016; Zhang and Sulzer, 2012) (Figure S2G). To test this we recorded single pulse-evoked DA release in dorsolateral striatum and applied increasing concentrations of the D2 receptor agonist quinpirole to construct a dose-response curve. Application of quinpirole decreased peak-evoked [DA]_o_ to a similar extent in DA-Tsc1 WT and DA-Tsc1 KO slices (WT IC50=102.7 ± 0.7933 nM, KO IC50=115.9 ± 0.8080 nM, Figure 2E,F). Therefore, it is unlikely that changes in D2 autoreceptor occupancy or expression account for the reduction in evoked DA release upon loss of Tsc1.

Striatal DA release is heavily dependent on voltage-gated calcium channels and is thus sensitive to changes in extracellular calcium levels (Rice et al., 2011; Sulzer et al., 2016). To test whether decreased DA release in DA-Tsc1 KO mice resulted from changes in the calcium sensitivity or calcium coupling of release sites, we tested if increasing extracellular calcium to 5 mM could rescue release deficits. As expected, increasing extracellular calcium led to increased peak-evoked [DA]_o_ in both genotypes compared to normal aCSF containing 2.4 mM Ca^2+^ (see Figure 2C,G). However, DA release in DA-Tsc1 KO slices remained ~60% lower compared to DA-Tsc1 WT slices (Figure 2G,H). Given the inability of increased extracellular calcium to normalize DA release deficits in DA-Tsc1 KO slices, we conclude that calcium sensitivity or coupling are unlikely to be major contributors to the release deficits observed.

### DA release probability (Pr) is significantly reduced following *Tsc1* deletion

One of the cell-intrinsic features that controls peak-evoked [DA]_o_ is release probability (Pr) (Cragg, 2003; Rice and Cragg, 2004), and a significant reduction in evoked [DA]_o_ may be indicative of compromised Pr. To address this possibility, we examined DA Pr in paired DA-Tsc1 WT and DA-Tsc1 KO slices using two approaches. First, we compared the ratio of peak [DA]_o_ evoked by 4 stimuli at 100 Hz to that evoked by a single pulse (4p/1p ratio, Figure 2I). The 4p/1p ratio was significantly higher in DA-Tsc1 KO mice compared to WT across all conditions, including drug-free aCSF, nAChR blockade (DHßE), and 5 mM extracellular Ca^2+^ (Figure 2J). These results indicated reduced initial Pr, a conclusion that was supported by the increased frequency-response sensitivity observed in DA-Tsc1 KO mice for short trains of four pulses delivered at 5-100 Hz (Figure S2H). Enhanced frequency sensitivity and 4p/1p ratio are reflective of the changes in somatic excitability we observed (see Figure 1), where sufficiently large depolarization could drive high firing rates in DA-Tsc1 KO neurons. Given the fundamental importance of the contrast between low level tonic and high level phasic DA signals (Berke, 2018), it is therefore plausible that these two modes of DA signaling to downstream circuitry remain at least partially intact following loss of Tsc1.

We next examined the paired-pulse ratio of release events evoked by single stimuli delivered three seconds apart, in the presence of DHßE to remove cholinergic modulation and 5 mM Ca^2+^ to maximize release probability (Figure 2K). The second-to-first peak ratio was significantly higher in DA-Tsc1 KO slices compared to DA-Tsc1 WT slices (Figure 2L). This indicates reduced suppression of DA release with repeated stimulation, again pointing to a reduction in initial dopamine Pr with Tsc1 loss.

During the FCV experiments, we observed that clearance of [DA]_o_, as measured by the falling phase of the DA transient, was faster in DA-Tsc1 KO slices (Figure S2I). Termination of the DA signal in the striatum is primarily mediated by uptake of released neurotransmitter back into the axon by the dopamine active transporter (DAT) (Hoffman et al., 1998; Nirenberg et al., 1996). Since uptake is exponentially related to the substrate concentration and DAT expression can vary by striatal region (Rice and Cragg, 2008; Rice et al., 2011), we identified concentration- and striatal region-matched transients from DA-Tsc1 WT and DA-Tsc1 KO slices and examined their decay kinetics. Curve-fit analysis of these transients showed that DA reuptake was significantly faster in DA-Tsc1 KO animals (Figure S2J). Combined, these findings suggest that Tsc1 loss from DA neurons strongly impairs their ability to influence striatal targets through the release of DA.

### DA neurons survive, innervate striatum, and have increased DA production following loss of Tsc1

To test whether the reduction in evoked DA release was a result of decreased numbers of DA neurons in DA-Tsc1 KO mice, we performed cell counts of tdTomato-labeled neurons in the SNc and VTA (Figure 3A,B). At three months of age, the number of tdTomato-expressing neurons was not significantly different between littermate DA-Tsc1 WT and DA-Tsc1 KO mice (Figure 3C, p=0.4343, unpaired, two-tailed t test). Therefore, deletion of *Tsc1* around embryonic day 15, when the first recombination events are detectable in the midbrain (Bäckman et al., 2006), does not strongly impact the development or survival of DA neurons into young adulthood.

**Figure 3.**
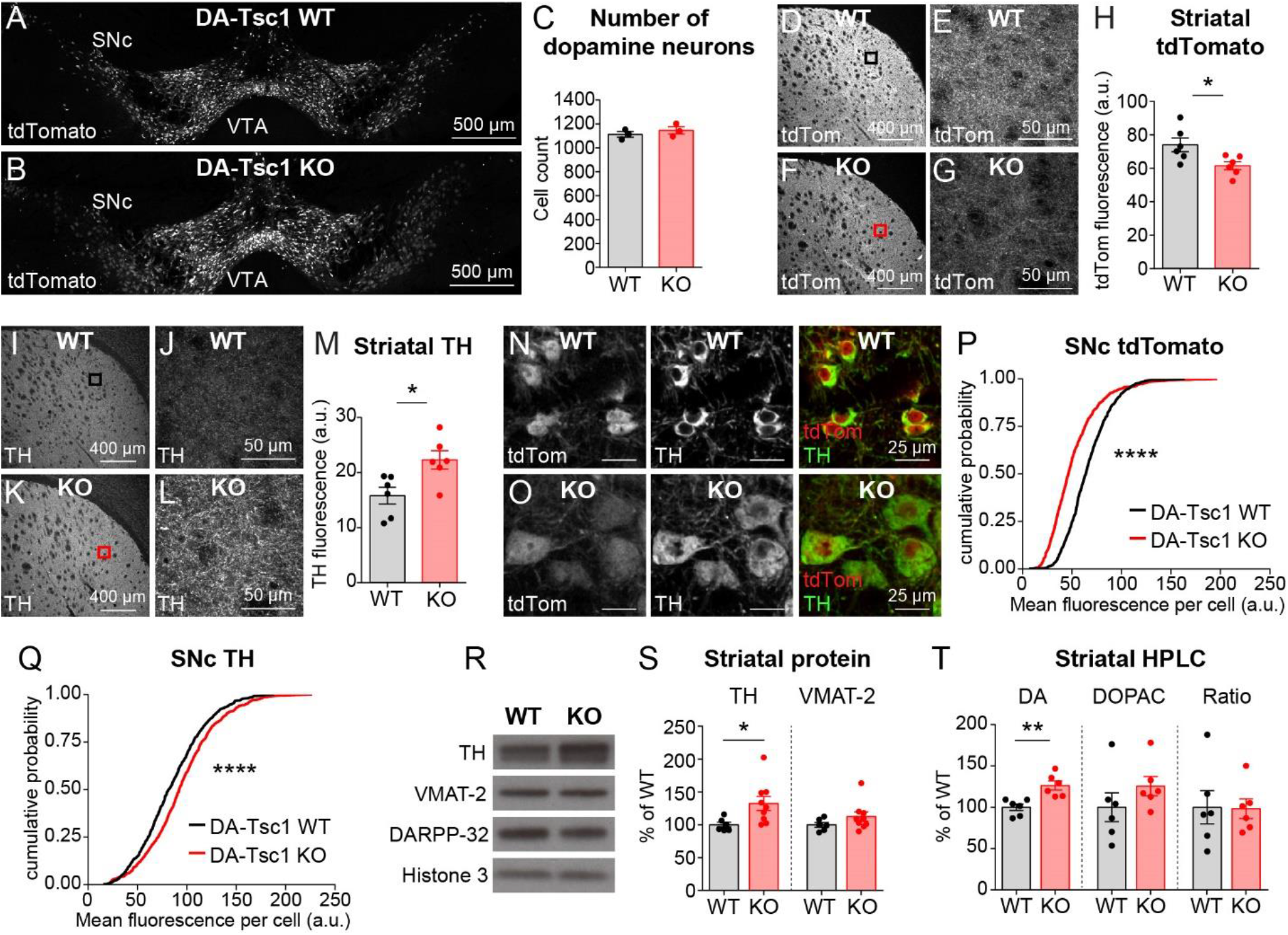
DA-Tsc1 KO neurons survive, innervate striatum, and synthesize more DA than controls. (A,B) Confocal images of coronal midbrain sections from *DAT^IRES^Cre^wt/+^;Ai9^wt/+^* mice homozygous for wild-type (DA-Tsc1 WT; A) or floxed *Tsc1* alleles (DA-Tsc1 KO; B). The tdTomato Cre reporter is expressed in DA neurons of the substantia nigra pars compacta (SNc) and ventral tegmental area (VTA). (C) Mean ± SEM number of tdTomato-positive DA neurons (including SNc and VTA) from anatomically matched coronal sections. Dots represent the mean cell count per mouse (9 sections were counted from 3 mice per genotype). p=0.4343, unpaired, two-tailed t test. (D-G) Low- (D,F) and high-magnification (E,G; from the boxed regions in panels D and F) confocal images of dorsolateral striatum from DA-Tsc1 WT (D,E) and DA-Tsc1 KO (F,G) mice showing tdTomato-positive DA axons. (H) Mean ± SEM tdTomato fluorescence intensity in dorsolateral striatum. Dots represent data from individual mice, n=6 mice per genotype (2 sections were analyzed per mouse). *, p=0.0265, unpaired, two-tailed t-test. (I-L) Low- (I,K) and high-magnification (J,L; from the boxed regions in I and K) confocal images of dorsolateral striatum from DA-Tsc1 WT (I,J) and DA-Tsc1 KO (K,L) mice showing tyrosine hydroxylase (TH) immunolabeling. (M) Mean ± SEM TH immunofluorescence intensity in dorsolateral striatum. Dots represent data from individual mice, n=6 mice per genotype (2 sections were analyzed per mouse). *, p=0.0173, unpaired, two-tailed t test. (N,O) Confocal images of DA-Tsc1 WT (N) and DA-Tsc1 KO (O) tdTomato-positive SNc DA neurons immunolabeled for TH. Right panels show merged images with tdTomato in red and TH in green. (P,Q) Cumulative probability plots of tdTomato (P) and TH (Q) levels in SNc DA neurons. Black lines show the distribution of values from DA-Tsc1 WT mice (n=759 neurons from 3 mice) and red lines show the distribution from DA-Tsc1 KO mice (n=855 neurons from 3 mice). ****, p<0.0001, Kolmogorov-Smirnov test. (R) Representative Western blot examples for TH and VMAT-2. The striatal-enriched protein DARPP-32 was used to normalize striatal tissue content and histone-3 was used as a loading control. (S) Mean ± SEM striatal protein content of TH and VMAT-2 expressed as a percent of WT. Dots represent average protein content per mouse, n=6 DA-Tsc1 WT mice and 9 DA-Tsc1 KO mice (2 striatum samples were analyzed per mouse). *, p=0.0169, unpaired, two-tailed t test. (T) Mean ± SEM striatal tissue content of DA and 3,4-dihydroxyphenylacetic acid (DOPAC) measured by high performance liquid chromatography (HPLC) with electrochemical detection. Data are expressed as a percent of WT. Ratio=DOPAC/DA ratio per mouse. Dots represent values from individual mice, n=6 mice per genotype (2 samples were averaged per mouse). **, p=0.0043, unpaired, two-tailed t test.

To test whether innervation of the striatum by DA axons might be impacted by loss of Tsc1, we analyzed tdTomato and TH expression in the dorsolateral striatum by immunofluorescence (Figure 3D-M). While tdTomato+ axons were present throughout the striatum in DA-Tsc1 KO mice, tdTomato fluorescence intensity was reduced compared to WT (Figure 3D-H). By contrast, levels of TH, the rate limiting enzyme necessary for DA synthesis, were significantly increased in DA-Tsc1 KO striatal axons (Figure 3I-M). To further explore these findings, we quantified tdTomato and TH levels in DA neuron cell bodies and found similarly decreased tdTomato and increased TH expression following loss of Tsc1 (Figures 3N-Q). Together, these results suggest that 1) the expression of tdTomato and TH are differentially regulated in the context of high mTORC1 signaling and 2) while dopaminergic axons still innervate the striatum in DA-Tsc1 KO mice, precise quantification of axon density is confounded by changes in axonal protein expression.

To investigate whether striatal axons in DA-Tsc1 KO mice have the capacity to synthesize and package DA, we harvested whole striata for western blot analysis and examined expression levels of TH and the vesicular monoamine transporter-2 (VMAT-2), which is responsible for loading DA into vesicles (Cartier et al., 2010). Consistent with the immunofluorescence results, we observed a significant increase in TH expression in the striatum of DA-Tsc1 KO mice compared to WT littermates, while VMAT-2 levels were unchanged (Fig 3R,S). These results indicate that DA-Tsc1 KO striatal axons have two of the key components necessary to synthesize and package DA into synaptic vesicles.

Higher TH levels suggest that DA-Tsc1 KO neurons may have increased capacity to synthesize DA. To examine this, we analyzed DA tissue content in the dorsal striatum using high-performance liquid chromatography (HPLC) with electrochemical detection. Consistent with high levels of TH, the total striatal tissue content of DA was significantly elevated DA-Tsc1 KO mice compared to WT (Figure 3T). However, levels of the primary DA metabolite DOPAC were not significantly affected (Figure 3T). Since the ratio of DOPAC to DA per mouse was unchanged by loss of Tsc1 (Figure 3T), we conclude that the high DA levels were due to increased DA synthesis rather than reduced turnover. The primary serotonin metabolite 5-hydroxyindoleacetic acid (5HIAA) could be detected in a subset of samples and was unchanged by *Tsc1* deletion (DA-Tsc1 KO=101.6% ± 15.25% of WT, p=0.966, unpaired, two-tailed t test), indicating selective alterations to the DA system.

### Loss of Tsc1 increases DA axon terminal size and reduces vesicle density

Our analyses showed a significant impairment in striatal DA release in DA-Tsc1 KO mice despite elevated striatal DA tissue content. To investigate whether structural changes to the axon terminals of DA neurons could be responsible for the release deficits, we performed electron microscopy (EM) analysis of DA terminals in the dorsolateral striatum identified by TH immunogold labeling (referred to as “profiles”, Figure 4A-D). DA neuron axons arborize extensively in the striatum, releasing DA at both synaptic and non-synaptic sites (Matsuda et al., 2009; Moss and Bolam, 2008; Rice and Cragg, 2008). We found that the majority of TH-expressing axon profiles were non-synaptic (Figure 4A,B), defined by the lack of a discernable synaptic membrane specialization. The percentage of synaptic profiles was similar between DA-Tsc1 WT and KO mice (11.25% and 10.25%, respectively) and both synaptic and non-synaptic axon profiles were included in the main analysis. Table S3 provides a separate analysis of synaptic profiles.

**Figure 4.**
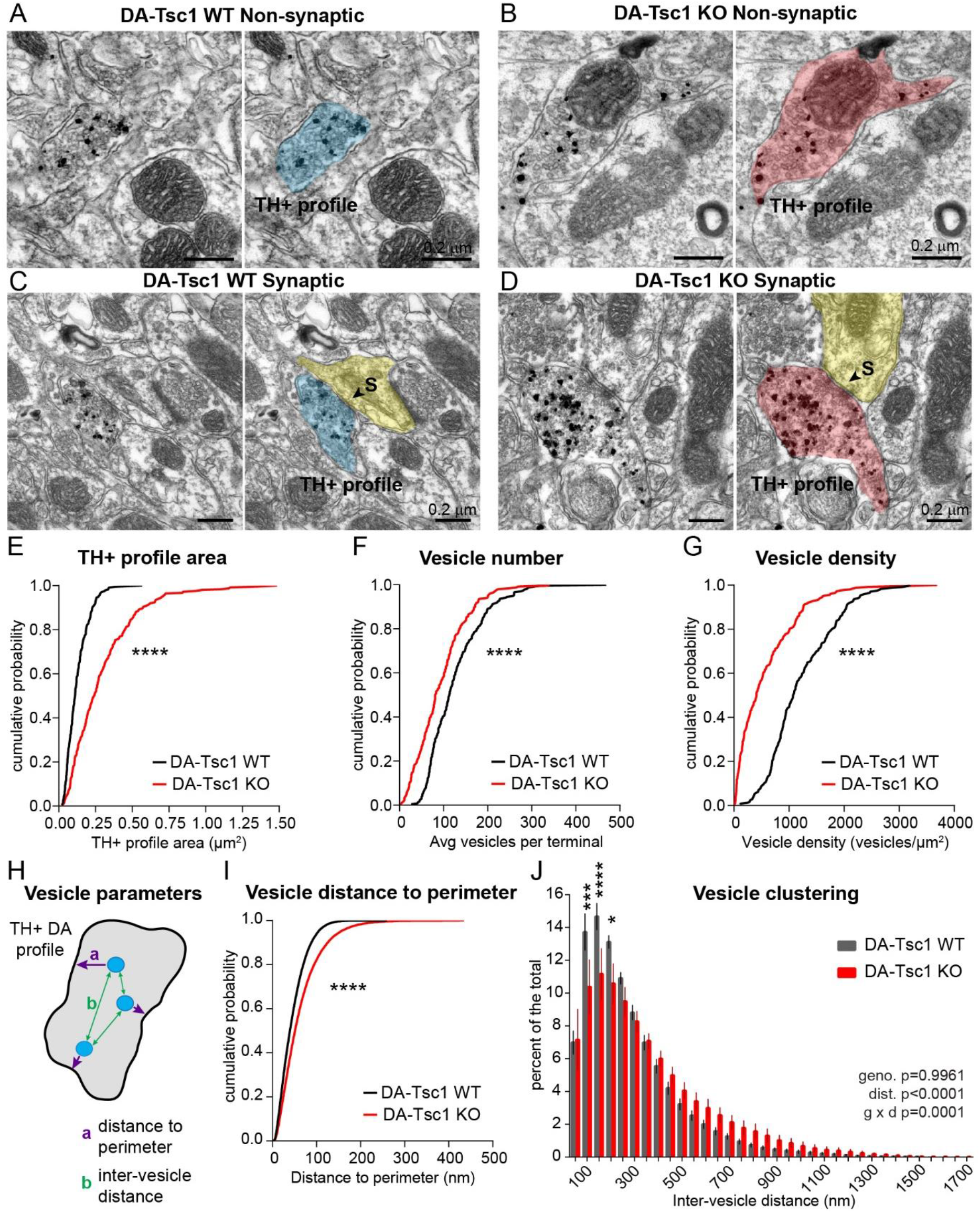
Loss of Tsc1 causes enlargement of DA axon terminals and reduced vesicle density and clustering. (A-D) Example electron micrographs (EM) of dopaminergic axon profiles in the dorsolateral striatum of DA-Tsc1 WT (A,C) and DA-Tsc1 KO (B,D) mice, identified by enriched immunogold labeling for tyrosine hydroxylase (TH). Right panels show TH+ axon profiles pseudocolored in blue (DA-Tsc1 WT) or red (DA-Tsc1 KO). A subset of axon profiles were observed to form synapses (“S”) (C-D), defined by the presence of a synaptic membrane specialization (arrowheads). The post-synaptic spine (C) or dendrite (D) is pseudocolored in yellow in the right panels. (E-G) Cumulative probability plots of TH+ axon profile area (E), vesicle number (F), and vesicle density (G) measured from EM of dorsolateral striatum.****, p<0.0001, Kolmogorov-Smirnov tests. (H) Schematic of vesicle position parameters measured within a TH+ axon profile. (I) Cumulative probability plot of vesicle distance to nearest point of the profile perimeter (i.e. plasma membrane). ****, p<0.0001, Kolmogorov-Smirnov test. (J) Histogram of mean ± SEM inter-vesicle distance in TH+ profiles in dorsolateral striatum binned at 50 nm. Two-way ANOVA p values are shown. *, p=0.0135, ***, p=0.0001, ****, p<0.0001, Sidak’s multiple comparisons test. For all panels, DA-Tsc1 WT: n=239 terminals (synaptic and non-synaptic) from 4 mice; DA-Tsc1 KO: n=252 terminals from 4 mice. See Table S3 for complete EM analysis.

Consistent with the somatodendritic hypertrophy we observed (see Figure 1), axonal profile area was significantly enlarged in DA-Tsc1 KO mice (Figure 4E and Table S3). We quantified the number of vesicles associated with each TH+ profile and found that vesicle number was significantly decreased in DA-Tsc1 KO profiles (Figure 4F). The combination of increased profile area, together with reduced vesicles per profile, led to a significant decrease in vesicle density DA-Tsc1 KO axons (Figure 4G). To determine the spatial arrangement of vesicles in each sampled axon, we measured the shortest distance of each vesicle to the profile perimeter (i.e. the plasma membrane), and the distance of each vesicle to every other vesicle within the profile (Figure 4H). We found that vesicle distance to the perimeter was significantly higher in DA-Tsc1 KO terminals compared to WT (Figure 4I). In addition, the average inter-vesicle distance was increased in DA-Tsc1 KO axons (DA-Tsc1 WT: 273.1 ± 0.14 nm, DA-Tsc1 KO: 303.7 ± 0.19 nm, p<0.0001, Kolmogorov-Smirnov test), indicating that vesicles were less clustered. When broken down into 50 nm bins, the histogram of inter-vesicle distances was shifted to the right for DA-Tsc1 KO terminals revealing significantly fewer vesicles within 100-200 nm of each other compared to WT terminals (Figure 4J). Thus, with reduced vesicle density and clustering, and vesicles further removed from the plasma membrane, the EM analysis of DA-Tsc1 KO axons revealed structural correlates of the observed deficits in DA release.

### Locomotor, affective, and social behaviors are normal while cognitive flexibility is reduced in DA-Tsc1 KO mice

DA is involved in a variety of behaviors, from movement to reward learning to cognition (Schultz, 2005; Steinberg et al., 2013; Wickens et al., 2007). To determine whether the marked perturbations in the functional properties of DA neurons affected the behavior of DA-Tsc1 KO mice, we performed a panel of behavior assays that are sensitive to dopaminergic function. Table S4 reports all behavior results and Table S5 shows behavioral analysis by sex.

To examine general exploratory behavior and locomotor activity, DA-Tsc1 KO mice and WT littermates were tested in the open field (Figure 5A). Deletion of *Tsc1* from DA neurons did not affect exploratory locomotor behavior, measured as the distance traveled in the first 10 minutes of the test (DA-Tsc1 WT: 23.44 ± 2.57 m, DA-Tsc1 KO: 22.62 ± 2.08 m, p=0.8046, unpaired, two-tailed t-test). The total distance traveled over 60 minutes and the mean speed were also similar between DA-Tsc1 WT and DA-Tsc1 KO mice (Figure 5B,C). We observed no changes in rearing behavior or time spent grooming between the two groups (Figure 5D,E). Therefore, despite a ~60% reduction in evoked DA release, locomotor activity was unaffected by DA neuron-specific loss of *Tsc1*.

**Figure 5.**
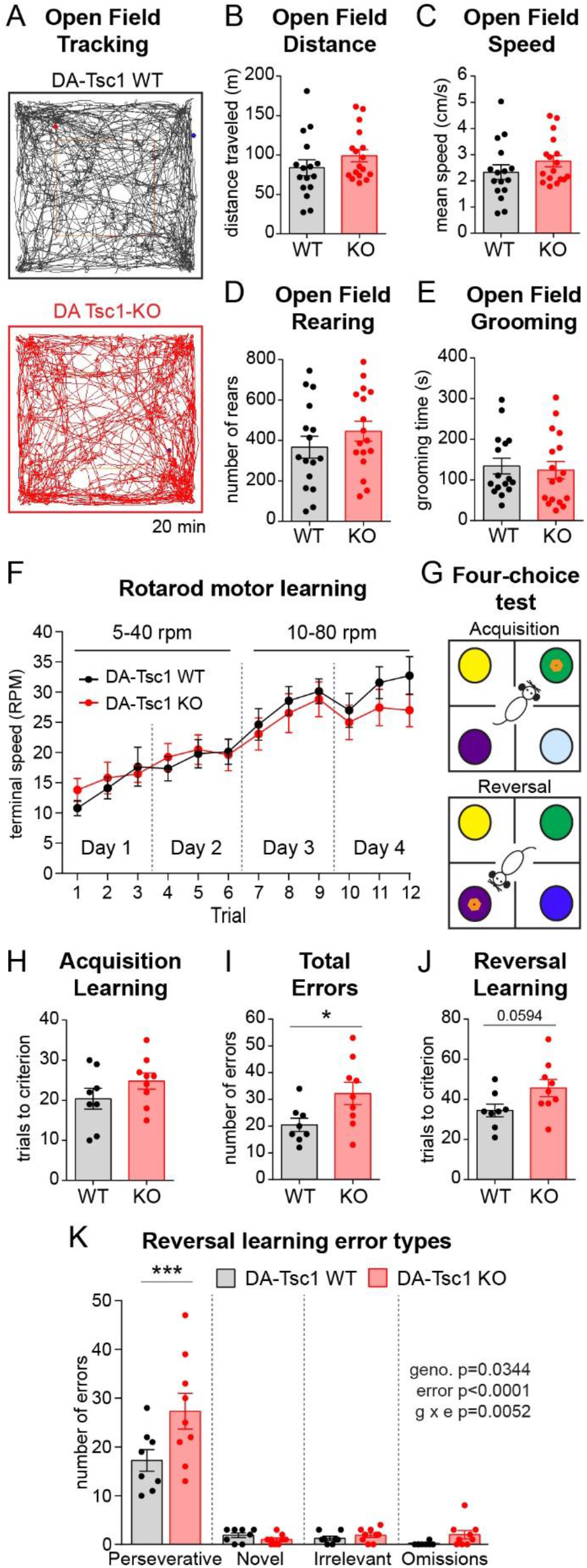
Loss of Tsc1 from DA neurons does not impact motor behavior but impairs cognitive flexibility. (A) Examples of automated tracking plots of mouse movement around the open field arena during first 20 minutes of the test. Movements of one mouse from each genotype are shown. (B-E) Quantification of open field behavior. Mean ± SEM total distance traveled over 60 minutes (B), mean speed (C), number of rears in 60 minutes (D), and time spent grooming during the first 20 minutes of the test (E). n=16 DA-Tsc1 WT mice and 17 DA-Tsc1 KO mice. Unpaired t tests revealed no significant differences. (F) Mean ± SEM terminal speed (speed of the rotarod when the mouse fell off) in revolutions per minute (rpm) for the accelerating rotarod test run over 4 days with 3 trials per day. n=12 DA-Tsc1-WT mice and 13 DA-Tsc1-KO mice. (G) Schematic of the 4-choice odor-based reversal task used to assess cognitive flexibility. On the first testing day (Acquisition), mice learn to associate one of four odors with a food reward (denoted by the orange ring). On the second testing day (Reversal), the mice have to learn a new odor-reward pairing and an unrewarded novel odor is introduced. (H-J) Mean ± SEM number of trials taken to reach criterion (8 out of 10 correct) during acquisition learning (H), total errors made during reversal learning (I), and trials to criterion during reversal learning (J). n=8 DA-Tsc1 WT mice and 9 DA-Tsc1 KO mice. *, p=0.0344, unpaired, two-tailed t test. (K) Analysis of different error types during reversal learning. Bars represent mean ± SEM number of perseverative, novel, irrelevant or omission errors for mice of each genotype. Two-way ANOVA p values are shown. **, p=0.0002, Sidak’s multiple comparisons test. For all panels, dots represent values for individual mice. See also Figure S3 and Tables S4 and S5 for all p values and behavior analysis by sex.

To test whether motor coordination or motor learning were altered in DA-Tsc1 KO mice, we performed the accelerating rotarod test (Rothwell et al., 2014). On days one and two of training, the acceleration ramped from 5 to 40 rpm over the course of five minutes and on days three and four, the acceleration ramped from 10 to 80 rpm. We found no differences between genotypes in either initial motor coordination (defined as the intercept of the line of best fit from trial 1 to 9; DA-Tsc1 WT 10.55 ± 1.19 rpm, DA-Tsc1 KO 13.37 ± 1.39 rpm, p=0.2904, Mann-Whitney test) or motor learning, measured as the slope of performance from the first to last trial for each mouse (DA-Tsc1 WT 1.92 ± 0.25, DA-Tsc1 KO 1.32 ± 0.19, p=0.0637, unpaired t test, Figure 5F). Together, our data indicate that these aspects of motor behavior are robust in the face of changes in the DA system caused by *Tsc1* deletion.

In addition to motor behavior, dopamine signaling modulates affective behaviors including anxiety and sociability (Manduca et al., 2016; Zweifel et al., 2011). We found no differences in measures of anxiety in the open field test and elevated plus maze between DA-Tsc1 WT and DA-Tsc1 KO mice (Figure S3A-H). Sociability was also normal in DA-Tsc1 KO mice as assessed by the three-chamber social approach test (Figure S3I-K).

The lack of motor deficits in DA-Tsc1 KO mice suggests that the motor system can compensate for developmental alterations in DA function, which is reflective of findings in Parkinson’s disease (PD) patients that motor impairments are only manifest when there is a >60-70% loss of DA terminals in the caudate-putamen (Cheng et al., 2010). However, cognitive dysfunction can be observed early in PD (Watson and Leverenz, 2010; Williams-Gray et al., 2007). In addition, cognitive flexibility, the ability to adapt behavior in response to a changing environment, is disrupted both in PD and in a range of neuropsychiatric conditions involving dopaminergic dysfunction (Cools et al., 2001; Klanker et al., 2013).

To examine associative learning and cognitive flexibility in DA-Tsc1 KO mice, we used the four-choice odor-based reversal task (Figure 5G) (Johnson et al., 2016), in which reversal learning is sensitive to changes in DA neuron function (Luo et al., 2016). Only male mice were used for this test, consistent with a prior study (Luo et al., 2016). In this test, mice learn to dig for a buried food reward in a pot of scented wood shavings. In the acquisition phase, mice learn an initial odor-reward pairing and the number of trials to reach criterion (8 out of 10 sequential trials correct) is a measure of discrimination learning. Following overnight consolidation, mice undergo a reversal learning phase in which the reward contingency is changed and a previously unrewarded odor is rewarded.

We found that DA-Tsc1 KO mice exhibited normal acquisition learning on the first day of testing, reaching criterion in a similar number of trials as littermate controls (Figure 5H). By contrast, when the rewarded odor was changed, DA-Tsc1 KO mice made significantly more total errors (Figure 5I) and tended to require more trials to learn the new odor-reward pairing (Figure 5J). For both genotypes, the majority of errors were perseverative, meaning that the mice continued to choose the initially learned odor even when it was no longer rewarded. However, DA-Tsc1 KO mice made significantly more perseverative errors than littermate controls (Figure 5K). The number of novel errors (choosing a newly introduced odor), irrelevant errors (choosing an odor that was never rewarded), or omissions was not significantly different between genotypes (Figure 5K). Thus, DA-Tsc1 KO mice exhibited a selective deficit in cognitive flexibility, wherein they were slower to update their behavioral strategy in the face of changed environmental conditions.

### Genetic reduction of *Rptor* constrains mTORC1 signaling in DA-Tsc1 KO neurons

Our data show that loss of Tsc1 from DA neurons strongly alters their structure, impairs their ability to release DA, and leads to reduced cognitive flexibility. To test whether these phenotypes could be prevented by suppression of mTORC1 signaling, we conditionally deleted one or two copies of the *Rptor* gene encoding the mTOR binding protein Raptor from DA neurons in DA-Tsc1 KO mice (Figures 6A,B and S4A). Raptor is an obligate member of mTORC1 and deletion of *Rptor* leads to selective mTORC1 loss of function (Kim et al., 2002; Sengupta et al., 2010). We found that homozygous deletion of *Rptor* from DA-Tsc1 KO mice (*Tsc1^fl/fl^;Rptor^fl/fl^;DAT^IRES^Cre^wt/+^*, referred to as “DA-Tsc1-KO/Rptor-KO”) strongly suppressed mTORC1 activity in DA neurons, as assessed by p-S6 levels (Figure S4B-F). Consistent with low mTORC1 signaling, DA neuron soma size was significantly smaller in DA-Tsc1-KO/Rptor-KO mice, and TH levels were reduced (Figure S4B-H). However, complete loss of Raptor was not able to rescue DA release deficits in DA-Tsc1 KO mice (Figure S4I-L), demonstrating that overactivation or complete suppression of mTORC1 signaling are similarly detrimental to striatal DA transmission.

**Figure 6.**
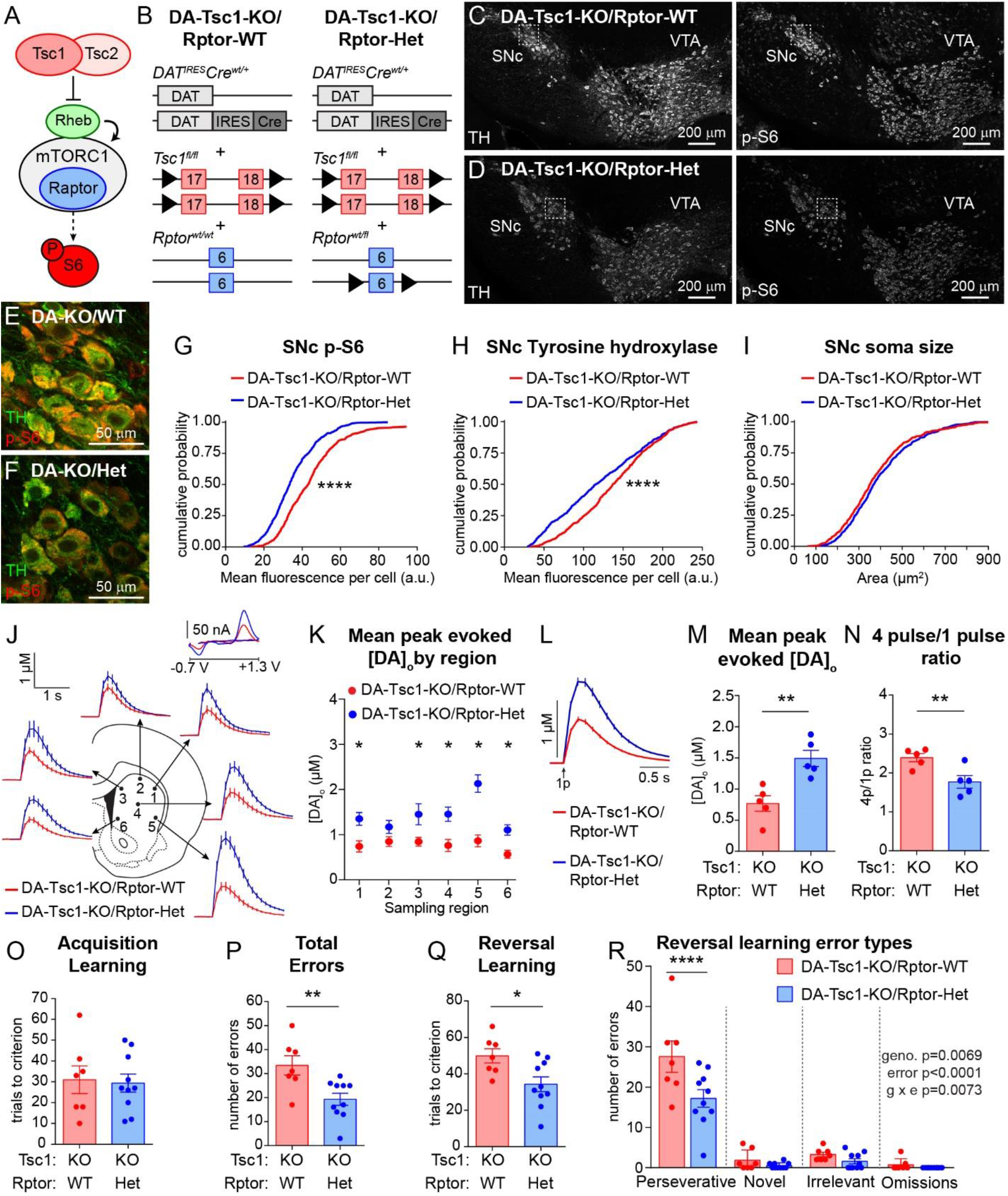
Heterozygous deletion of *Rptor* prevents DA release deficits and improves cognitive flexibility in DA-Tsc1 KO mice. (A) Simplified mTORC1 signaling schematic showing the Tsc1/2 heterodimer as a negative regulator of the small GTP-ase Rheb, which directly promotes mTORC1 kinase activity. Raptor is an essential component of mTORC1. mTORC1 regulates the phosphorylation of ribosomal protein S6 via p70S6 kinase (represented by the dashed arrow). (B) Schematic of the genetic strategy to selectively delete *Tsc1* and reduce *Rptor* by one copy in DA neurons. (C-F) Confocal images of coronal midbrain sections from DA-Tsc1-KO/Rptor-WT (C,E) and DA-Tsc1-KO/Rptor-Het (D,F) mice. Sections were labeled with antibodies against tyrosine hydroxylase (TH) and phosphorylated S6 (p-S6, Ser240/244). Panels E and F show higher magnification merged images of the boxed regions in C and D. SNc=substantia nigra pars compacta, VTA=ventral tegmental area. (G-I) Cumulative probability plots of SNc DA neuron p-S6 levels (G), tyrosine hydroxylase (TH) levels (H), and soma area (I). Red lines show the distributions of values from DA-Tsc1-KO/Rptor-WT mice (n=528 neurons from 3 mice) and blue lines show distributions for DA-Tsc1-KO/Rptor-Het mice (n=567 neurons from 3 mice). ****, p<0.0001, Kolmogorov-Smirnov tests. (J) Mean extracellular DA release ([DA]_o_) ± SEM versus time evoked by single-pulse electrical stimuli from different striatal subregions. Traces show an average of 16-20 transients per recording site from 5 mice per genotype. 1-dorsolateral striatum, 2-dorsocentral striatum, 3-dorsomedial striatum, 4-central striatum, 5-ventrolateral striatum, 6-ventromedial striatum. Inset, typical cyclic voltammograms show characteristic DA waveform. (K) Mean peak single pulse-evoked [DA]_o_ ± SEM by striatal region (numbers correspond to the numbered sites in panel A). n=16-20 transients per recording site from 5 mice per genotype. *, p_1_=0.0002; *, p_3_=0.0.0150; *, p_4_=0.0011; *, p_5_<0.0001; *, p_6_=0.0005, paired t tests. (L) Mean single pulse-evoked [DA]_o_ ± SEM versus time for all dorsal striatum sites recorded in normal aCSF. Average of 112 transients across 6 recording sites per genotype from 5 mice. (M) Mean peak single pulse-evoked [DA]_o_ ± SEM averaged across all dorsal striatum sites (sites #1-6 in panels A and B). Dots represent average evoked [DA]_o_ per mouse in normal aCSF. n=5 mice per genotype. **, p=0.0087, paired t test. (N) Mean ± SEM ratio of peak [DA]_o_ evoked by a 4 pulse train to single pulse stimulation in dorsolateral striatum in normal aCSF. Dots represent the average 4p/1p ratio per mouse. n=5 mice per genotype. **, p=0.0062, paired t test. (O-Q) Quantification of the mean ± SEM number of trials taken to reach criterion (8 out of 10 correct) during acquisition learning (O), total errors made during reversal learning (P), and trials to criterion during reversal learning (Q) in the 4-choice test. n=7 DA-Tsc1-KO/Rptor-WT mice and 10 DA-Tsc1-KO/Rptor-Het mice. *, p=0.0183; **, p=0.0069, unpaired t tests. (R) Analysis of different error types during reversal learning. Bars represent mean ± SEM number of perseverative, novel, irrelevant or omission errors for mice of each genotype. Two-way ANOVA p values are shown. ****, p<0.0001, Sidak’s multiple comparisons test. For all panels, dots represent values for individual mice. See also Figure S4 and Table S4 for all p values for behavior tests.

We next tested whether reduction, rather than complete suppression, of mTORC1 signaling could prevent phenotypes in DA-Tsc1 KO mice. To do this we compared DA-Tsc1 KO mice that were either wild-type (*Tsc1^fl/fl^;Rptor^wt/wt^;DAT^IRES^Cre^wt/+^*, “DA-Tsc1-KO/Rptor-WT”) or heterozygous for the conditional *Rptor* allele (*Tsc1^fl/fl^;Rptor^wt/fl^;DAV^IRES^Cre^wt/+^*, “DA-Tsc1-KO/Rptor-Het”) (Figure 6B). Consistent with a reduction in mTORC1 signaling, we found that p-S6 levels in SNc neurons were significantly reduced in DA-Tsc1-KO/Rptor-Het mice compared to DA-Tsc1-KO/Rptor-WT (Figure 6C-G). Elevated TH levels in DA-Tsc1-KO/Rptor-WT SNc neurons were also reduced by heterozygous deletion of *Rptor* (Figure 6C-F and H). Despite reductions in p-S6 and TH, DA neurons in DA-Tsc1-KO/Rptor-Het mice did not have smaller somata (Figure 6C-F and I). Therefore, heterozygous *Rptor* gene deletion constrained mTORC1 signaling in DA neurons in the context of *Tsc1* loss; however, this partial reduction of mTORC1 activity was not sufficient to prevent somatic hypertrophy.

### Heterozygous *Rptor* deletion prevents DA release deficits in DA-Tsc1 KO mice and increases cognitive flexibility

To test whether the functional deficits in DA release in DA-Tsc1 KO mice could be prevented by genetic reduction of mTORC1 signaling, we performed FCV experiments in paired DA-Tsc1-KO/Rptor-WT and DA-Tsc1-KO/Rptor-Het striatal slices. We found that DA-Tsc1-KO/Rptor-Het slices had significantly higher evoked [DA]_o_ throughout the dorsal striatum compared to DA-Tsc1-KO/Rptor-WT slices (Figure 6J,K). On average, evoked DA levels were two-fold higher in the dorsal striatum of DA-Tsc1-KO/Rptor-Het mice compared to DA-Tsc1-KO/Rptor-WT (Figure 6L,M). We compared the 4p/1p ratio between the two experimental groups to estimate DA release probability, and found that DA-Tsc1-KO/Rptor-Het slices had significantly lower 4p/1p ratio compared to DA-Tsc1-KO/Rptor-WT (Figure 6N). Therefore, heterozygous *Rptor* deletion alleviated DA release deficits, likely via a significant increase in dopamine Pr.

Given the functional improvements in DA release that we observed in DA-Tsc1-KO/Rptor-Het mice, we tested whether these mice had improved cognitive flexibility, as assessed by their performance in the four-choice reversal learning task. We observed no significant differences between DA-Tsc1-KO/Rptor-Het and DA-Tsc1-KO/Rptor-WT male littermate mice during acquisition learning (Figure 6O). However, during reversal learning, DA-Tsc1-KO/Rptor-Het animals made fewer total errors (Figure 6P), reached criterion in significantly fewer trials (Figure 6Q), and made fewer perseverative errors (Figure 6R). Together, these data demonstrate that cell type-specific genetic reduction of mTORC1 signaling in DA-Tsc1-KO mice can improve evoked DA release and counter reductions in cognitive flexibility.

## Discussion

Dopaminergic circuits play a central role in the cognitive functions and behaviors that are impacted in mTOR-related neurodevelopmental disorders. Here we used genetic mouse models to investigate how deregulation of mTORC1 signaling affects the functional properties of DA neurons. We find that DA neuron-specific loss of Tsc1, a key negative regulator of mTORC1, leads to constitutive activation of mTORC1 signaling in DA neurons, profoundly affecting their somatodendritic and axonal architecture, intrinsic excitability, and ability to release DA in the dorsal striatum. Mice lacking Tsc1 selectively in DA neurons exhibit reduced cognitive flexibility in the absence of changes to motor function, consistent with patients with mutations in *TSC1* or *TSC2* (de Vries et al., 2015). We show that DA release deficits and cognitive inflexibility in DA-Tsc1 KO mice can be prevented by genetic reduction of mTORC1 signaling. Together these data provide insight into the cellular origins of the behavioral manifestations associated with deregulated mTORC1 signaling, and suggest a potential locus for intervention.

### mTOR signaling in DA neurons

The impact of mTOR signaling on DA neuron physiology has primarily been studied in the context of addiction. Specifically, drugs of abuse have been shown to activate mTORC1 signaling in midbrain DA neurons (Collo et al., 2012, 2013; Dayas et al., 2012; Neasta et al., 2014; Wu et al., 2011), and pharmacologic or genetic inhibition of mTOR changes synaptic function in VTA neurons and attenuates the behavioral effects of drugs of abuse (Dayas et al., 2012; Liu et al., 2018b; Neasta et al., 2014; Wu et al., 2011). mTORC2 signaling has also been implicated in reward tolerance to opiate drugs via changes in VTA neuron size and excitability (Mazei-Robison et al., 2011).

mTORC1 signaling has also been studied in the context of PD, where mTOR activation has been shown to acutely promote DA neuron survival, possibly via increased protein synthesis at the site of injury and/or inhibition of autophagic processes (Cheng et al., 2011; Diaz-Ruiz et al., 2009; Domanskyi et al., 2011; Kim et al., 2012; Lan et al., 2017; Malagelada et al., 2010; Xu et al., 2014; Zhou et al., 2015). Here, we build upon this literature and show, for the first time, that chronic developmental activation of mTORC1 signaling arising from Tsc1 loss leads to altered DA neuron structure at multiple levels, inducing somatic, dendritic, and axonal hypertrophy. These morphological changes are associated with decreased intrinsic excitability and impaired DA release. Together with prior studies, this work establishes mTOR as an essential regulator of DA neuron output, which influences multiple aspects of DA neuron function, including membrane excitability and pre- and post-synaptic properties.

### DA neuron function requires balanced mTORC1 signaling

We found that chronic developmental activation of mTORC1 signaling profoundly affected DA neuron structure and significantly impaired neurotransmitter release. We attempted to prevent these alterations by inhibiting mTORC1 signaling via deletion of the gene encoding the obligate mTORC1 protein Raptor in DA-Tsc1 KO mice. However, we found identical deficits in evoked DA release in DA-Tsc1-KO/Rptor-KO mice as with *Tsc1* deletion alone. Release deficits persisted despite complete reversal of *Tsc1* KO-induced somatic hypertrophy. The failure of complete mTORC1 suppression to prevent release deficits is consistent with prior reports showing that acute treatment of striatal slices with the mTOR inhibitor rapamycin or Cre virus-mediated deletion of *mTOR* from VTA neurons both decrease evoked DA release (Hernandez et al., 2012; Liu et al., 2018b). By contrast, we found that heterozygous loss of *Rptor* in DA-Tsc1 KO mice, while not sufficient to prevent somatic hypertrophy, significantly improved DA release and countered reduced cognitive flexibility. Together, these results indicate that tight regulation of mTORC1 signaling is essential for DA neuron output as too much or too little mTORC1 signaling is similarly detrimental to DA release. In addition, our findings show that changes in DA neuron soma size can be decoupled from alterations in axonal DA release

### Mechanisms of DA release deficits in DA-Tsc1 KO mice

DA neurons have specialized intrinsic and extrinsic mechanisms to control DA release (Liu et al., 2018a; Sulzer et al., 2016). We investigated several potential mechanisms for the DA release deficits in DA-Tsc1 KO mice. We first ruled out DA neuron death, striatal denervation, and non-cell-autonomous effects on the striatal cholinergic system as potential explanations. We also excluded changes in D2 autoreceptor expression or occupancy. In addition, we found that increasing extracellular calcium was not sufficient to normalize release, suggesting that mechanisms other than (or in addition to) calcium sensitivity or coupling to the release machinery were involved. Importantly, we showed that the axons of DA neurons in DA-Tsc1 KO mice had key molecular machinery to synthesize and package DA into vesicles and, in fact, had higher TH levels, as reported previously for DA neurons with increased mTORC1 signaling due to *Pten* deletion (Diaz-Ruiz et al., 2009; Domanskyi et al., 2011). High TH levels implied greater capacity to synthesize DA. Indeed, we found that the striatal tissue content of DA was significantly increased in DA-Tsc1 KO mice compared to controls. This may have been driven by increased mTORC1-dependent protein synthesis of TH. Alternatively, TH expression could have been increased independent of mTORC1, as a homeostatic mechanism to compensate for decreased DA release.

The most plausible mechanisms for impaired DA release in DA-Tsc1 KO mice were ultrastructural changes in DA neuron axons. Specifically, we found that Tsc1 loss highly enlarged TH+ axon terminals in the dorsal striatum. Axon profiles in DA-Tsc1 KO mice also had fewer vesicles. The net result of these changes was a marked reduction in vesicle density. In addition, the vesicles were, on average, further removed from the plasma membrane and were less tightly clustered together. Given the stringent structural requirements for efficient vesicle release, for example vesicle proximity to docking sites (Park et al., 2012), it is likely that the observed ultrastructural changes in DA-Tsc1 KO axons led to reduced Pr and impaired DA release. Further, in a mouse model of PD overexpressing alpha-synuclein, abnormal clustering of vesicles alone caused a 30% reduction in evoked striatal DA release (Janezic et al., 2013).

A recent study demonstrated that a subset of striatal DA terminals have active zone protein scaffolds, which contain RIM as an essential component and are required for DA release (Liu et al., 2018a). It will be interesting for future studies to explore the potential contribution of changes in active zone proteins to mTOR-dependent effects on DA release.

### Behavioral consequences of mTOR-related dopaminergic dysfunction

A notable finding of our study is that despite severe impairments in dorsal striatal DA release, DA-Tsc1 KO mice did not exhibit gross motor impairments. This was assessed by multiple parameters in the open field (total distance traveled, average speed, rearing) and accelerating rotarod test, which measures motor coordination and learning. While initially surprising, the lack of motor deficits is reminiscent of observations in PD where motor symptoms are only evident after 30-40% of SNc neurons have degenerated and striatal DA axons are reduced by 60-70% (Cheng et al., 2010). However, cognitive deficits can be observed in PD patients even at early stages of the disease, suggesting that cognitive functions may be particularly vulnerable to changes in DA (Watson and Leverenz, 2010; Williams-Gray et al., 2007). In addition, in DA-Tsc1 KO mice, *Tsc1* is deleted embryonically, therefore, the basal ganglia circuit develops in the context of reduced dopaminergic output and compensatory mechanisms are likely engaged in dopaminoceptive neurons.

We also found that anxiety and social approach behavior were unchanged in DA-Tsc1 KO mice. These behaviors are modulated by mesolimbic DA projections originating from the VTA (Manduca et al., 2016; Zweifel et al., 2011). The lack of change in these behaviors is consistent with the fact that DA release in the nucleus accumbens (a major projection target of VTA neurons) was not significantly altered by *Tsc1* deletion. It is also possible that manipulation of mTORC1 signaling in DA neurons alone is not sufficient to strongly affect social behavior.

This contrasts with cerebellar Purkinje cell-specific deletion of *Tsc1*, which does impair social behavior in a rapamycin-sensitive manner (Tsai et al., 2012). Together, these studies reveal that different cell types are differentially vulnerable to changes in mTORC1 signaling and that distinct cell types and circuits are likely responsible for specific aspects of mTOR-related disorders.

Cognitive flexibility, defined as the ability to adapt behavior in the face of changing environmental demands, is commonly affected in psychiatric disorders, in particular ASD and ADHD (D’Cruz et al., 2013; Dajani and Uddin, 2015; Sergeant et al., 2002). TSC patients frequently present with impairments in executive control processes, which include cognitive flexibility (Prather and de Vries, 2004; de Vries et al., 2015). Here, we found that DA neuron-specific deletion of *Tsc1* was sufficient to reduce cognitive flexibility in a reversal-learning task. This impairment was specific to reversal learning as initial discrimination learning was intact. Notably, during reversal learning, DA-Tsc1 KO mice made more perseverative errors, in which they continued to return to the previously learned odor even though it was no longer rewarded. Perseverative errors reflect an inability to flexibly adapt and “update” a previously learned association. Such a deficit may contribute to the inflexible behaviors often observed in patients with TSC (de Vries et al., 2015).

Our findings are consistent with prior reports showing that brain-wide heterozygous deletion of *Tsc2* or disruption of the translational regulator eIF4E, which increases protein synthesis, impair reversal but not acquisition learning in water maze tasks (Potter et al., 2013; Santini et al., 2013). Our work suggests that these types of impairments may be attributable to dopaminergic dysfunction, consistent with the involvement of DA in cognitive flexibility in rodents and humans (Darvas and Palmiter, 2011; Klanker et al., 2013). Our findings are also in agreement with studies showing that moderate reductions in dorsal striatal DA levels in rats (as achieved by toxin injections) can impair cognitive flexibility without changing motor behavior (Grospe et al., 2018; O’Neill and Brown, 2007).

Taken together, our work demonstrates that developmental deregulation of mTORC1 signaling profoundly alters key functional properties of DA neurons. These changes lead to reduced dopaminergic output and a selective deficit in cognitive flexibility. Our findings have implications for cognitive dysfunction in neurodevelopmental disorders associated with altered mTORC1 signaling and suggest that these phenotypes may be driven by dopaminergic dysfunction.

## Acknowledgments

We thank the members of the Bateup lab for their feedback on this project. We thank Dr. Linda Wilbrecht and members of the Wilbrecht lab for assistance with the four-choice reversal learning task. We thank Dr. Bernardo Sabatini for the original idea to investigate Tsc1 loss in dopamine neurons. We thank Ben Micklem for his technical support in maintaining the electron microscope facility at the MRC Brain Network Dynamics Unit. This work was supported by a NARSAD Young Investigator Grant from the Brain & Behavior Research Foundation (#25073) and a Sloan Research Fellowship in Neuroscience (#FR-2015-65789) to H.S.B. P.K. was supported by a post-doctoral research grant from the Tuberous Sclerosis Alliance (2015 TS Alliance Research Grants Program #381490). The work of N.M.D., L.C. and P.J.M. was funded by the Medical Research Council of the United Kingdom (award MC_UU_12024/2) and the Wellcome Trust (Investigator Award 101821 to P.J.M.). The work of S.T. was supported by Parkinson’s UK Monument Trust Discovery Award (award J-0901).

## Author Contributions

Conceptualization, P.K. and H.S.B.; Methodology, P.K., N.M.D., S.T., P.J.M. and H.S.B.; Formal Analysis, P.K., N.M.D., A.A.M., K.A., S.T., and H.S.B.; Investigation, P.K., N.M.D., A.A.M., K.A., L.C., and S.T.; Writing – Original Draft, P.K. and H.S.B.; Writing – Review & Editing, P.K., N.M.D., S.T., P.J.M., and H.S.B.; Visualization, P.K., N.M.D., and H.S.B.; Supervision, P.J.M. and H.S.B.; Funding Acquisition, P.K., H.S.B. and P.J.M..

## STAR Methods

### CONTACT FOR REAGENT AND RESOURCE SHARING

Further information and requests for resources and reagents should be directed to and will be fulfilled by the Lead Contact, Dr. Helen Bateup

### EXPERIMENTAL MODEL AND SUBJECT DETAILS

#### Mice

All animal procedures were carried out in accordance with protocols approved by the University of California, Berkeley Institutional Animal Care and Use Committee (IACUC). The ages of the animals used are indicated in the Method Details for each experiment. Unless otherwise indicated, mice of both sexes were used. Mice were an outbred strain on mixed genetic background as determined by the Charles River Mouse 384 SNP Complete Background Analysis Panel, with average percent heterozygosity of 22.9% (13.1%-33.7% range). Mice were housed with same sex littermates in groups of 5-6 animals per cage and kept on a regular 12 hr light/dark cycle (lights on at 7am), with ad libitum access to food and water. Animals in behavior experiments were housed in a facility with a reverse 12 hr light/dark cycle (lights off at 9am). Mice were switched to the reverse light cycle at least two weeks prior to testing and were tested during the dark phase.

##### Table of transgenic mouse lines

| Mouse strain | Description | Reference/Source |
| --- | --- | --- |
| DAT<sup>IRESc<sup>re</sup></sup> (referred to as DAT<sup>IRESCre</sup>) | Cre recombinase knocked in to the *Slc6a3* (DAT) locus | (Bäckman et al., 2006); Dr. Bernardo Sabatini lab |
| *Tsc1*<sup>fl/fl</sup> | Conditional *Tsc1* knock-out with loxP sites flanking exons 17 and 18 of the *Tsc1* gene | (Kwiatkowski et al., 2002); Dr. Bernardo Sabatini lab |
| *Rptor*<sup>fl/fl</sup> | Conditional *Rptor* knock-out with loxP sites flanking exon 6 of the *Rptor* gene | (Sengupta et al., 2010); Dr. Bernardo Sabatini lab |
| Ai9 | Cre reporter allele with CAG promoter lox-STOP-lox tdTomato cassette in the *Rosa26* locus | (Madisen et al., 2010); Jackson Laboratories stock #007909 |

##### Breeding strategy

To generate dopamine neuron-specific Tsc1 knock-out mice, *Tsc1^fl/fl^* mice were bred to *DAT^IRES^Cre^wt/+^* mice. To generate experimental animals, mice heterozygous for floxed *Tsc1* and *DAT^IRES^Cre* were crossed (*Tsc1^wt/fl^;DAT^IRES^Cre^wt/+^* x *Tsc1^wt/fl^;DAT^IRES^Cre^wt/+^*). Experimental mice were heterozygous for Cre and either homozygous wild-type for Tsc1 (*Tsc1^wt/wt^;DAT^IRES^Cre^wt/+^*, referred to as DA-Tsc1 WT) or homozygous floxed for Tsc1 (*Tsc1^fl/fl^; DAT^IRES^Cre^wt/+^*, referred to as DA-Tsc1 KO).

For electrophysiology experiments, *Tsc1^fl/fl^;DAT^IRES^Cre^wt/+^* mice were bred to the Ai9 tdTomato Cre-reporter line. *Tsc1^wt/fl^;DAT^IRES^Cre^wt/+^;Ai9^wt/+^* x *Tsc1^wt/fl^;DAT^IRES^Cre^wt/+^;Ai9^wt/+^* crosses were used to generate experimental mice. Mice used for experiments were either heterozygous or homozygous for the Ai9 transgene.

For rescue experiments, *Tsc1^fl/fl^;DAT^IRES^Cre^wt/+^* mice were crossed with *Rptor^fl/fl^* mice. To generate experimental animals, a *Tsc1^fl/fl^;Rptor^wt/fl^;DAT^IRES^Cre^wt/+^* x *Tsc1^fl/fl^;Rptor^wt/fl^;DAT^IRES^Cre^wt/+^* breeding strategy was used. The resulting offspring were all homozygous floxed for *Tsc1* and either wild-type, heterozygous, or homozygous floxed for *Rptor*.

##### Table of transgenic mice used for experimentation

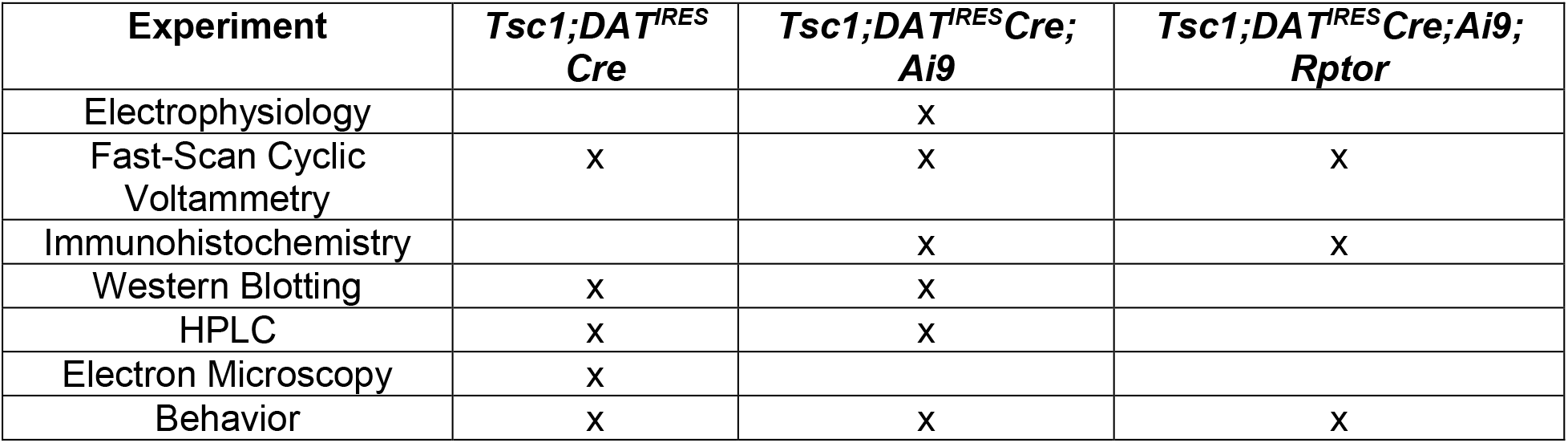

#### METHOD DETAILS

##### Electrophysiology

Male and female adult mice (P56-P80) were deeply anesthetized by isoflurane, transcardially perfused with ice cold high Mg^2+^ ACSF using a peristaltic pump (Instech) and decapitated. 275 μm thick coronal midbrain slices were prepared on a vibratome (Leica VT1000 S) in ice cold high Mg^2+^ ACSF containing in mM: 85 NaCl, 25 NaHCO_3_, 2.5 KCl, 1.25 NaH_2_PO_4_, 0.5 CaCl_2_, 7 MgCl_2_, 10 glucose, and 65 sucrose. Slices were recovered for 15 minutes at 34°C followed by at least 50 minutes at room temperature (RT) in ACSF containing in mM: 130 NaCl, 25 NaHCO_3_, 2.5 KCl, 1.25 NaH_2_PO_4_, 2 CaCl_2_, 2 MgCl_2_, and 10 glucose. All solutions were continuously bubbled with 95%O2 and 5% CO_2_.

Recordings were performed at 32°C in the presence of AMPA, NMDA and GABA_A_ synaptic blockers (10 μM GYKI 52466; 10 μM CPP; 50 μM picrotoxin, final concentration, all from Tocris), with a bath perfusion rate of 2 ml/min. Dopaminergic neurons in the SNc and VTA were identified by Ai9 tdTomato fluorescence. For whole cell recordings, 2.5-6 mΩ borosilicate glass pipettes (Sutter Instruments: BF150-86-7.5) were filled with a potassium-based internal solution containing (in mM): 135 KMeSO_3_, 5 KCl, 5 HEPES, 4 Mg-ATP, 0.3 Na-GTP, 10 phospho-creatine, 1 EGTA, and 4mg/ml neurobiotin (Vector laboratories #SP-1120). Recordings were obtained using a MultiClamp 700B amplifier (Molecular Devices) and ScanImage software. Passive membrane properties were recorded in voltage clamp with the membrane held at −70 mV. Positive current steps (2 s, 25-600 pA) were applied to generate an input-output curve from a baseline membrane potential of −70 mV as maintained by hyperpolarizing current injection. For whole-cell recordings, series resistance was <30 MΩ and liquid junction potential was not corrected.

##### Biotin-Filled Neuron Reconstruction

DA neurons were filled with neurobiotin-containing internal solution (4 mg/ml) during whole-cell recordings. Slices were fixed in 4% paraformaldehyde solution (Electron Microscopy Sciences: 15713) in 1x PBS for 24-48 h at 4°C. With continuous gentle shaking, slices were washed in 1x PBS 3 x 5 minutes and incubated with BlockAid blocking solution (Life Tech: B10710) for 1 h at RT. Primary antibodies against tyrosine hydroxylase (1:1000, Immunostar: 22941) and streptavidin-Alexa Flour 488 conjugate (1:750, Invitrogen: S11223) were applied overnight at 4°C in 1x PBS containing 0.25% Trinton-X-100 (PBS-Tx). The following day, slices were washed in 1x PBS 3 x 5 minutes, and secondary Alexa-633 goat anti-mouse antibody (1:500, ThermoFisher: A21050) in PBS-Tx was applied for 1 h at RT. Slices were washed in cold 1x PBS 5 x 5 min, mounted on SuperFrost slides (VWR: 48311-703) with the neurobiotin-filled cell facing up, and coverslipped with either Prolong Gold antifade (Life Tech: P36935) or Vectashield hard-set (Vector Labs: H-1500) mounting media.

Neurobiotin-labelled cells were imaged on a Zeiss LSM 880 NLO AxioExaminer confocal with 20x/1.0 N.A. water immersion objective and 488 Argon laser using 1.53 μm steps to acquire a z-stack image spanning the entirety of the neurobiotin-filled cell body and dendritic arbor. 3D reconstruction of the cells was performed using IMARIS software (Bitplane) with automated filament tracing and manual editing. The mask generated by the automated filament tracing algorithm was continuously cross-referenced with the original z-stack image to ensure accuracy. Spurious segments created by the automated filament tracer due to background noise were removed, while processes with incomplete reconstruction were edited to incorporate missing segments. Sholl analysis of dendritic arborization was performed by quantifying the number of intersections of dendrites with concentric circles drawn at 1 μm steps, starting 5 μm from the center point of the soma.

##### Fast-Scan Cyclic Voltammetry (FCV)

DA release was monitored using fast-scan cyclic voltammetry (FCV) in acute coronal slices as described previously (Rice and Cragg, 2004; Threlfell et al., 2010, 2012). Paired site-sampling recordings were performed in one reference and one experimental animal in a sex- and age-matched mouse pair recorded on a given day. The genotype order of tissue prep was counterbalanced between experiments. Male and female mice (P56-P95) were deeply anesthetized by isoflurane and decapitated. 275 μm thick coronal striatal slices were prepared on a vibratome (Leica VT1000 S) in ice cold high Mg^2+^ ACSF containing in mM: 85 NaCl, 25 NaHCO_3_, 2.5 KCl, 1.25 NaH_2_PO_4_, 0.5 CaCl_2_, 7 MgCl_2_, 10 glucose, and 65 sucrose. Slices were recovered for 1 h at RT and were recorded from in ACSF containing in mM: 130 NaCl, 25 NaHCO_3_, 2.5 KCl, 1.25 NaH_2_PO_4_, 2 CaCl_2_, 2 MgCl_2_, and 10 glucose. All solutions were continuously saturated with 95%O2 and 5% CO_2_. Slices between +1.5 mm and +0.5 mm from Bregma containing dorsal striatum and nucleus accumbens (ventral striatum) were used for experimentation (Paxinos and Franklin, 2008).

In the recording chamber, slices were maintained at 32°C with a superfusion rate of 1.2-1.4 ml/min. Extracellular DA concentration ([DA]_o_) was monitored with FCV at carbon-fiber microelectrodes (CFMs) using a Millar voltammeter (Julian Millar, Barts and the London School of Medicine and Dentistry). CFMs were fabricated in-house from epoxy-free carbon fiber ~7 μm in diameter (Goodfellow Cambridge Ltd) encased in glass capillary (Harvard Apparatus: GC200F-10) pulled to form a seal with the fiber and cut to final tip length of 70–120 μm. The CFM was positioned ~ 100 μm below the tissue surface at a 45 degree angle. A triangular waveform was applied to the carbon fiber scanning from −0.7 V to +1.3 V and back, against a Ag/AgCl reference electrode at a rate of 800 V/s. Evoked DA transients were sampled at 8 Hz, and data were acquired at 50 kHz using AxoScope 10.5 (Molecular Devices). Oxidation currents evoked by electrical stimulation were converted to [DA]_o_ from post-experimental calibrations of the CFMs. Recorded FCV signals were identified as DA by comparing oxidation (+0.6 V) and reduction (−0.2 V) potential peaks from experimental voltammograms with currents recorded during calibration with 2 μM dopamine dissolved in ACSF and all drug-containing media used in a given experiment.

##### Electrical stimulation and drug application

Following 30 min slice equilibration in the recording chamber, DA release was evoked, using square wave pulses (0.6 mA pulse amplitude, 2 ms pulse duration) controlled by a Master-8 pulse stimulator or Isoflex stimulus isolator (A.M.P.I., Jerusalem, Israel) delivered out of phase with voltammetric scans. A concentric bipolar stimulating electrode (FHC: CBAEC75) used for electrical stimulation was positioned on the slice surface first with minimal tissue disturbance followed by CFM inserted 100 μm away from the bipolar electrode. Three stimulation paradigms were used: single stimulation pulse, paired single pulses delivered 3 s apart, and brief 4-pulse stimulation trains delivered at 5, 10, 25, 40 or 100 Hz. Stimuli were delivered every 2.5 min.

For site sampling experiments, three stimulations were delivered at a given site before moving to the corresponding site in the slice from the paired mouse. Stimulations were delivered in the following order: single pulse, pulse train of 4 pulses at 100 Hz, single pulse.

For quinpirole experiments, after reaching a steady baseline release with single pulses in control ACSF, increasing doses of quinpirole (1 nM-10 μM, Tocris) were bath-applied for ~ 10 minutes each, with steady-state release achieved for at least four consecutive stimulations before progression to the next highest drug concentration.

For current-amplitude and frequency-response recordings, recordings were performed from a single site per slice. After reaching a steady baseline release with single pulses, each stimulation condition was repeated 3 times at each frequency/current amplitude in a pseudorandom order.

##### Immunohistochemistry

Male and female mice (P75-P95) were deeply anesthetized by isoflurane and exsanguinated by transcardial perfusion with ice cold 1x PBS (~5-7 ml), followed by 4% paraformaldehyde (PFA) solution (Electron Microscopy Sciences: 15713) in 1x PBS (~5-10 ml) using a peristaltic pump (Instech). The brains were removed and post-fixed by immersion in 4% PFA in 1x PBS overnight at 4°C. Brains were then suspended in 30% sucrose in 0.1 M PB solution for cryoprotection. After brains descended to the bottom of the vial (typically 24-28 h), 30 μm coronal sections of the midbrain and striatum were cut on a freezing microtome (American Optical AO 860), collected into serial wells and stored at 4°C in 1x PBS containing 0.02% (w/v) sodium azide (NaN_3_; Sigma Aldrich). All mice used for histology experiments were heterozygous for the Ai9 tdTomato allele.

Free-floating brain sections for immunohistochemistry were batch processed to include matched control and experimental animals from a given mouse line. With gentle shaking, sections were washed 3 x 5 min in 1x PBS followed by 1 h incubation at RT with BlockAid blocking solution (Life Tech: B10710). Primary antibodies were applied at 4°C in 1x PBS containing 0.25% Trinton-X-100 (PBS-Tx) for 48-72 h. Sections were washed with cold PBS 3 x 5 min, incubated for 1 h at RT with secondary antibodies in PBS-Tx, washed in cold PBS 5 x 5 min, mounted on SuperFrost slides (VWR: 48311-703), and coverslipped with Vectashield hard-set (Vector Labs: H-1500) mounting media. For DA neuron count experiments, slices underwent no additional processing apart from 1x PBS wash 5 x 5 min. Experimenters were blind to animals’ genotype throughout tissue processing and data analysis.

The following primary antibodies were used: tyrosine hydroxylase (1:1000, Immunostar: 22941) and phospho-S6 (Ser240/244) (1:1000, Cell Signaling: 5364S). The following secondary antibodies were used: Alexa-488 goat anti-mouse (1:500, Thermo Fisher: A-11001) and Alexa-633 goat anti-rabbit (1:500, Thermo Fisher: A-11034).

##### Confocal Microscopy

Images of 30 μm sections processed for immunohistochemistry were acquired using a Zeiss LSM 710 AxioObserver confocal microscope fitted with a motorized XY-stage for tile scanning. A 20x/0.8 N.A. air objective was used to generate tile scans (424×424 μm per tile) of one hemisphere (5×3 grid) or the entire midbrain (10×3 grid). 405 nm, 488 nm, 561 nm and 633 nm laser were used. Z-stack images captured the entire thickness of the slice at 1.10-1.25 μm steps. Laser power settings and acquisition parameters were kept constant for all experimental conditions.

##### High-Performance Liquid Chromatography (HPLC) with Electrochemical Detection

Tissue DA content was measured by HPLC with electrochemical detection in tissue punches from dorsal striatum. Male and female mice (P60-P75) were deeply anesthetized by isoflurane and decapitated. 300 μm thick coronal slices of striatum were prepared on a vibratome (Leica VT1000 S) in ice cold high Mg^2+^ ACSF containing in mM: 85 NaCl, 25 NaHCO_3_, 2.5 KCl, 1.25 NaH_2_PO_4_, 0.5 CaCl_2_, 7 MgCl_2_, 10 glucose, and 65 sucrose. Slices were recovered for 1 hour at RT in ACSF containing in mM: 130 NaCl, 25 NaHCO_3_, 2.5 KCl, 1.25 NaH_2_PO_4_, 2 CaCl_2_, 2 MgCl_2_, and 10 glucose. All solutions were continuously bubbled with 95%O2 and 5% CO_2_. Following slice recovery, tissue punches from the dorsal (2.5 mm diameter) striatum from two brain slices per animal were taken and stored at −80°C in 200 μl 0.1 M HClO4. On the day of analysis, samples were thawed, homogenized and centrifuged at 16,000 x g for 15 min at 4°C. The supernatant was analyzed for DA, 3,4-Dihydroxyphenylacetic acid (DOPAC), and 5-Hydroxyindoleacetic acid (5HIAA) content using HPLC with electrochemical detection. Analytes were separated using a 4.6 x 150 mm Microsorb C18 reverse-phase column (Varian or Agilent) and detected using a Decade II SDS electrochemical detector with a Glassy carbon working electrode (Antec Leyden) set at +0.7 V with respect to a Ag/AgCl reference electrode. The mobile phase consisted of 13% methanol (v/v), 0.12 M NaH2PO4, 0.5 mM OSA, 0.8 mM EDTA, pH 4.8, and the flow rate was fixed at 1 ml/min. Analyte measurements were normalized to tissue punch volume (pmol/mm^3^). HPLC analysis was repeated in two independent experiments.

##### Western Blotting

Male and female mice (P60-P90) were deeply anesthetized by isoflurane and decapitated. Bilateral striata were rapidly dissected on ice, flash-frozen in liquid nitrogen and stored at −80°C. On the day of analysis, frozen samples were sonicated until homogenized (QSonica Q55) in 500 μl lysis buffer containing 1% SDS in 1x PBS with Halt phosphatase inhibitor cocktail (Fisher: PI78420) and Complete mini EDTA-free protease inhibitor cocktail (Roche: 4693159001). Sample homogenates were then boiled on a heat block at 95°C for 10 min, allowed to cool down to RT, and total protein content was determined by BCA assay (Fisher: PI23227). Following BCA assay, protein homogenates were mixed with 4x Laemmli sample buffer (Bio-Rad: 161-0747).10-15 μg of protein were loaded onto 4–15% Criterion TGX gels (Bio-Rad: 5671084). Proteins were transferred to PVDF membrane (BioRad: 1620177) at 4°C overnight using the Bio-Rad Criterion Blotter (12 V constant voltage). The membranes were blocked in 5% milk in 1x TBS with 1% Tween (TBS-T) for 1h at RT, and incubated with primary antibodies diluted in 5% milk in TBS-T overnight at 4°C. The following day, after 3 x 10 min washes with TBS-T, the membranes were incubated with HRP-conjugated secondary antibodies for 1h at RT. Following 6 x 10 min washes, the membranes were incubated with chemiluminesence substrate (Perkin-Elmer: NEL105001EA) for 1 min and exposed to GE Amersham Hyperfilm ECL (VWR: 95017-661). Membranes were stripped with re-blot plus strong solution (Millipore: 2504) to re-blot on subsequent days. The following primary antibodies were used: mouse anti-tyrosine hydroxylase (1:3000, Immunostar: 22941); mouse anti-DARPP-32 (1:3000, gift from Dr. Paul Greengard’s Lab); mouse anti-Histone-3 (1:1500, Cell Signaling: 96C10); rabbit anti-VMAT2 (1:1400, Alomone Labs: AMT-006). Secondary antibodies were goat anti-rabbit HRP (1:5000, Bio-Rad: 170-5046) and goat anti-mouse HRP (1:5000, Bio-Rad: 170-5047). Each western blotting experiment was repeated with at least two independent sets of samples. Experimenters were blind to genotype throughout WB processing and data analysis

##### Electron Microscopy (EM)

###### Sample preparation

Male and female mice (P75-P85) were deeply anesthetized by isoflurane, exsanguinated by transcardial perfusion with ice cold 0.1 M PB (pH 7.4), followed by fixation with 0.1% glutaraldehyde (Sigma Aldrich: G5882) and 4% paraformaldehyde (Electron Microscopy Sciences: 15713) solution in 0.1 M PB. Perfusions were performed using a peristaltic pump (Instech). The brains were removed and immersed in the fixative solution for an additional 12 h at 4°C. Brains were then stored in 0.1 M PB with 0.02% (w/v) NaN_3_. Coronal sections (50 μm) were cut using a vibrating blade microtome (Leica VT1000 S). All sections were washed 3 x 5 min in 1x PBS, placed into cryoprotectant (0.05 M PB, 25% sucrose, 10% glycerol) for a minimum of 2 h and then freeze-thawed three times in liquid nitrogen to increase penetration of the reagents. Sections were washed thoroughly and blocked with 10% normal goat serum in 1x PBS (Vector Laboratories: S-1000) for 1 h followed by overnight incubation in primary rabbit anti-tyrosine hydroxylase antibody (1:1000, Chemicon: #AB152) at RT. Sections were washed 3 x 5 min in 1x PBS and incubated in a gold-conjugated goat-anti-rabbit secondary antibody (1:400, 1.4 nm Nanogold, Nanoprobes: #2003) for a minimum of 4 h at RT. TH-positive axons were revealed with silver intensification of the conjugated gold particles. Silver reagent (1 ml; Nanoprobes: HQ Silver kit, prepared according to manufacturer’s instructions) was added to each section and allowed to react for 4.5 min in the dark, washed 3 x 5 min in acetate buffer (0.1 M sodium acetate 3-hydrate, pH 7.0–7.5) and then 1x PBS for 5 min. All sections were then washed 3 x 5 min in 0.1 M PB. The sections were post-fixed in 1% osmium tetroxide in 0.1 M PB (OsO_4_; TAAB) for 12 min. After washing 2 x 5 min in 0.1 M PB, sections were dehydrated in an ascending series of ethanol dilutions: 2 x 10 min in 50% ethanol; 45-60 min in 70% ethanol, which included 1% uranyl acetate (TAAB); 10 min in 95% ethanol; and 2 x 10 min in absolute ethanol. Sections were washed 2 x 10 min in propylene oxide (Sigma Aldrich), placed into resin (Durcupan ACM; Fluka), and left overnight (~15 h) at RT. The resin was then warmed to reduce its viscosity, and sections were placed on microscope slides, coverslipped, and the resin cured at 65°C for ~70 h (Doig et al., 2010, 2014). Experimenters were blind to genotype throughout tissue processing and data analysis.

###### Electron microscopy

All sections were examined under a light microscope and areas from the dorsolateral striatum were cut from the slide, glued to the top of a resin block, and trimmed with razor blades. For each mouse, striatal regions from two sections across the rostro-caudal plane were examined. Serial sections, ~50 nm thick, were cut using an ultramicrotome (Leica EM UC6), collected on pioloform-coated, single-slot copper grids (Agar Scientific), and lead-stained with Reynold’s lead citrate to improve contrast for electron-microscopic examination. For each region (block), ultrathin sections from at least two grids were examined. A Philips CM100 electron microscope was used to examine the sections. Analyses of pre-embedded immunogold sections were performed at a minimum of 5 μm from the tissue–resin border (i.e., section surface). The maximum distance from the tissue–resin border examined was determined by the penetration of the gold conjugated antibody together with the angle at which the tissue–resin was sectioned, and was therefore variable. TH-positive structures were systematically analyzed in one of the serial sections on an electron-microscopic grid. At a magnification at which it is not possible to clearly visualize synapses, an area was chosen at random, the magnification was then increased and the first structure positively labeled for TH was digitally recorded (GatanCCD UltraScan US1000 camera; Gatan). TH-positive structures were identified and imaged, in this way, continuing systematically in straight lines across the section, keeping the identified TH-positive structure central within the image frame. For immunogold-labeled structures, the criterion for an immunopositive structure was five or more silver-intensified immunogold particles. The process was continued within the same ultrathin section until 15 TH-positive structures were identified and imaged, and was then repeated using a different ultrathin section, on a different grid, from the same block. This was repeated for the second block until a minimum of 60 TH-immunopositive structures were identified and imaged per animal (491 dopaminergic profiles in total, n=4 mice per genotype). Symmetric synapses (Gray’s type II) were identified by the presence of presynaptic and postsynaptic membrane specializations, a widened synaptic cleft and cleft material. Any TH-positive structures seen to be forming such symmetric synapses were imaged (Anwar et al., 2011; Janezic et al., 2013).

##### Behavioral Tests

Behavior testing was carried out in the dark phase of the light cycle under red lights, except the rotarod and four-choice reversal tests, which were carried out in the dark phase under white lights. Mice were habituated to the behavior room for at least 30 min prior to testing and covered by a black-out curtain. Mice were given at least one day in between different tests. All behavior equipment was cleaned between each trial/mouse with 70% ethanol, and additionally rinsed in diluted soap followed by water at the end of the day. If male and female mice were to be tested on the same day, male mice were run in behavior first, returned to the husbandry room, after which all equipment was thoroughly cleaned, and female mice were brought in for habituation. Behavioral tests were performed with P60-80 male and female mice except for the four-choice test for which P70-130 male animals were used. The experimenter was blind to genotype throughout the testing and scoring procedures.

###### Open Field Test

Exploratory behavior in a novel environment and general locomotor activity were assessed by a 60 min session in an open field chamber (40 cm L x 40 cm W x 34 cm H) made of transparent plexiglas. Horizontal photobeams to measure rearing were positioned at 9 cm of height. The mouse was placed in the bottom right hand corner of the arena and behavior was recorded using an overhead camera and analyzed using the ANY-maze (Stoelting Co) behavior tracking software. An observer manually scored scratching and grooming behavior during the first 20 minutes of the test.

###### Rotarod Test

The accelerating rotarod test was used to examine motor coordination, balance and motor learning. Mice were run on a rotarod apparatus (Ugo Basile: 47650) for 4 consecutive days. Three trials were completed per day with a 5 min break between trials. The rotarod was accelerated from 5-40 revolutions per minute (rpm) over 300 s for trials 1-6 (days 1 and 2), and from 10-80 rpm over 300 s for trials 7-12 (days 3 and 4). On the first testing day, mice were acclimated to the apparatus by being placed on the rotarod rotating at a constant 5 rpm for 60 s and returned to their home cage for 5 minutes prior to starting trial 1. Latency to fall, or to rotate off the top of the rotarod barrel, was measured by the rotarod stop-trigger timer.

###### Four-Choice Odor-Based Reversal Task

The four-choice odor-based reversal task (Johnson et al., 2016) was used to assess learning and cognitive flexibility in adult male mice. Animals were food restricted for 5-6 days in total with unrestricted access to drinking water, and maintained at 85-90% of ad lib feeding body weight. Food was given at the end of the day once testing was completed. Food restriction began 24 h before pre-training.

The four-choice test was performed in a custom-made square box (30.5 cm L × 30.5 cm W × 23 cm H) constructed of clear acrylic. Four internal walls 7.6 cm wide partially divided the arena into four quadrants. A 15.2 cm diameter removable cylinder fit in the center of the maze and was lowered between trials (after a digging response) to isolate the mouse from the rest of the maze. Odor stimuli were presented in white ceramic pots measuring 7.3 cm diameter and 4.5 cm deep. Pots were sham baited with a cereal reward, either a Honey Nut Cheerio (General Mills, Minneapolis, MN) or Fruit Loop cereal (Kellogg’s) secured underneath a mesh screen at the bottom. The apparatus was cleaned with 2.5 % acetic acid followed by water between mice. Pots were cleaned with 70% ethanol followed by water. The apparatus was cleaned with diluted soap and water at the end of each testing day.

On the first habituation day of pre-training (day 1), animals were allowed to freely explore the testing arena for 30 minutes and consume small pieces of Cheerios placed inside pots positioned in each of the four corners. On the second shaping day of pre-training (day 2), mice learned to dig to find cereal pieces buried in unscented coarse pine wood shavings (Harts Mountain Corporation, Secaucus, NJ). On the shaping day, animals who failed to consume Cheerio pieces were switched to Fruit Loops, which they consumed successfully. A single pot was used and increasing amounts of unscented wood shavings were used to cover each subsequent cereal reward. The quadrant containing the pot was alternated on each trial and all quadrants were rewarded equally. Trials were untimed and consisted of (in order): two trials with no shavings, two trials with a dusting of shavings, two trials with the pot a quarter full, two trials with the pot half full, and four trials with the cereal piece completely buried by shavings.

The mouse was manually returned to the center cylinder between trials.

On the testing days for odor discrimination (day 3) and reversal (day 4), wood shavings were scented on the day of testing. Anise extract (McCormick, Hunt Valley, MD) was used undiluted at 0.02 ml/g of shavings. Clove, litsea, and eucalyptus oils (San Francisco Massage Supply Co., San Francisco, CA) were diluted 1:10 in mineral oil and mixed at 0.02 ml/g of shavings. Thymol (“thyme”; Alfa Aesar) was diluted 1:20 in 50% ethanol and mixed at 0.01 ml/g of shavings. During the discrimination phase (day 3), mice had to discriminate between four pots with four different odors and learn which one contained a buried food reward (*indicates the rewarded odor in the table below). Each trial began with the mouse confined to the start cylinder, once the cylinder was lifted, timing began and the mouse could freely explore the arena until it chose to dig in a pot. Digging was defined as purposefully moving the shavings with both front paws. A trial was terminated if no choice was made within 3 min and recorded as omission. Criterion was met when the animal completed 8 out of 10 consecutive trials correctly. The reversal phase of the task was completed the following day (day 4). Mice first performed the task with the same rewarded odor as the discrimination day to ensure they learned and remembered the task. After reaching criterion on recall (8 out of 10 consecutive trials correct), the rewarded odor was switched. Perseverative errors were choices to dig in the previously rewarded odor that was no longer rewarded. Novel errors were choices to dig in the pot with the newly introduced odor for reversal testing. Irrelevant errors were choices to dig in the pot that had never been rewarded. Omissions were trials in which the mouse failed to make a digging choice within three minutes from the start of the trial. Total errors are the sum of perseverative, irrelevant, novel and omission errors. Criterion was met when the mouse completed 8 out of 10 consecutive trials correctly.

|  | **Odor 1** | **Odor 2** | **Odor 3** | **Odor 4** |
| --- | --- | --- | --- | --- |
| Acquisition | *anise | clove | litsea | thyme |
| Reversal | anise | *clove | litsea | eucalyptus |

###### Elevated Plus Maze Test

Exploratory behavior in a novel environment and generalized anxiety were assessed with elevated plus maze test (Walf and Frye, 2007). The maze consisted of four arms 30 cm long and 5 cm wide, with two open arms and two arms enclosed by 16 cm tall walls painted black. The apparatus was custom made out of Plexiglas plastic painted black and attached to sturdy Plexiglas plastic legs, which elevated it 39 cm off the floor. At the start of the test, the mouse was placed in the center square of the apparatus facing an open arm and allowed to freely explore for 5 min. Behavior was recorded using a video camera positioned directly above the center of the plus maze and analyzed using the ANY-maze (Stoelting Co) behavior tracking software. An entry into a closed or open arm was counted when the mouse had all four paws in the arm.

###### Three Chamber Social Approach Test

Social approach behavior was assessed using the three chamber social approach test (Yang et al., 2011). The testing apparatus was made of clear Plexiglas (60 cm L× 40 cm W × 22 cm H, Stoelting). Two dividing walls had doorways allowing access to three chambers. Age and sex matched C57BL/6J mice were used as novel mice and were habituated to placement inside the wire cups (8 cm diameter, 11 cm tall) prior to testing in either three 10-minute or two 15-minute sessions. C57BL/6J mice that displayed persistent agitation, including climbing or biting behavior inside the wire cups, were not used for testing.

The test mouse was placed in the center chamber of the apparatus and allowed to explore and habituate to all three empty chambers for 10 min. The mouse was then placed in the center chamber again for 5 min with access doors closed. During this time, a novel C57BL/6J mouse was placed under a wire cup in one of the side chambers and an empty wire cup was placed on the opposite side (novel object). The chamber doors separating the three chambers were then removed to open access to both left and right chambers. The mouse was allowed to explore the three chambers for 10 min, and allowed to freely interact with the novel object or with the stranger mouse inside the wire cup. The location of the stranger mouse and the novel object were alternated between mice. Behavior was recorded using a video camera positioned directly above the center of the apparatus and analyzed using the ANY-maze (Stoelting Co) behavior tracking software. An observer blind to genotype manually scored time spent sniffing the novel mouse and the novel object during the test. Sniffing was defined as when the mouse’s nose was perpendicular to the vertical plane and less than 1 cm away from the wire cup. If the mouse reared on the wire cup, climbed on the wire cup, or bit the wire cup, it was not scored as sniffing. Mice showing a strong side bias during the habituation phase of the test were excluded. Controls were *DAT^IRES^Cre* negative littermates of DA-Tsc1 KO mice. Comparisons were made within genotype.

### QUANTIFICATION AND STATISTICAL ANALYSES

Whenever possible, quantification and analyses were performed blind to genotype. All statistical analyses and graphing were performed using GraphPad Prism 6 software. All datasets were first analyzed using D’Agostino and Pearson omnibus normality test, and then parametric or non-parametric two-tailed statistical tests were employed accordingly to determine significance. Significance was set as *, p < 0.05; **, p < 0.01; ***, p < 0.001; ****, p < 0.0001 unless otherwise indicated in the figure legend. P values were corrected for multiple comparisons. The statistical tests used for each type of experiment are indicated below. Statistical details for each experiment are reported in the figure legends.

#### Electrophysiology Data

Data were analyzed in Igor Pro (Wavemetrics) using custom scripts. Evoked action potential properties were measured from the first AP in the train of evoked APs. AP threshold was calculated by taking the second derivative of the action potential. Passive properties and action potential properties were analyzed for significance using unpaired t-tests. Excitability curves were analyzed for significance using a two-way ANOVA with Sidak’s multiple comparison test.

#### Dopamine neuron reconstruction

Morphological parameters, including Sholl intersections and total dendritic length, were extracted using the IMARIS (Bitplane) statistics feature from masks. Total dendrite length was analyzed using an unpaired t-test and Sholl intersections using a repeated measures two-way ANOVA.

#### Voltammetry Data

Data were pre-processed using AxoScope 10.5 (Molecular Devices) software and analyzed using custom written Excel macros and scripts (Prof S. Cragg and Dr K. Jennings, Oxford University). Drug data were normalized to control data, and frequency data were normalized to single pulses, before collating across experiments. We included a minimum of three release events for each stimulus or condition at each individual recording site, unless otherwise stated.

For all site sampling experiments (aCSF, DHβE, oxo-M, 5 mM Ca^2+^) single pulse and four pulses at 100 Hz-evoked DA concentrations were compared without normalization because control and experimental slices were recorded with the same CFM for every experimental pair.

A minimum of four matched mouse pairs were used in these experiments, with two slices per animal recorded from. Single pulse data includes two release events per site per slice, pulse train data includes one release event per site per slice. Peak evoked DA release levels were compared between genotypes using paired t-test tests. Quinpirole and DAT uptake kinetics were compared using curve-fit analyses. Current-amplitude and frequency-response data were compared using a two-way ANOVA with Sidak’s multiple comparisons test.

#### Protein Data

##### Immunohistochemistry

Analyses were performed blind to genotype. Quantification was performed on max-projected z-stack images. For soma size and p-S6 measurement, soma boundaries were traced manually using Image J (NIH) software in the tyrosine hydroxylase channel to include all clearly identifiable single cells in the SNc or VTA (minimum 100 cells for each area in each slice). Soma cross-sectional area and fluorescence intensity for the p-S6 channel were extracted automatically in Image J. Cumulative distributions were analyzed using the Kolmogorov-Smirnov test.

For DA neuron counts, every clearly identifiable tdTomato+ cell in both hemispheres was manually counted using the Cell Counter Image J (NIH) plug-in, and analyzed using an unpaired t test. Three anatomically matched sections in the rostro-caudal plane per mouse were counted in three age- and sex-matched mouse pairs.

##### Western Blotting

Western blot analysis was performed blind to genotype. Bands were quantified by densitometry using Image J (NIH) software. Proteins were normalized to DARPP-32 and expressed as percentage of control within a given experiment. TH data were analyzed using an unpaired t-test with Welch’s correction and VMAT-2 data were analyzed using a Mann-Whitney test.

##### HPLC

Analysis was performed blind to genotype. Data were normalized and expressed as a percent of control within a given experiment before collating across experiments (samples were collected and analyzed in two batches). Data were analyzed for statistical significance using unpaired t-test.

#### Electron Microscopy Data

Analysis was performed blind to genotype. Digital images were analyzed using Image J (NIH) and the Image J plug-ins PointDensity and PointDensitySyn (https://liu.se/medfak/forskning/larsson-max/software?l=en; Larsson et al., 2015). Images were adjusted for contrast and brightness using Photoshop (version CS5; Adobe Systems). For analysis of TH-positive structures, the central TH-positive structure within the image frame was analyzed by tracing the perimeter; and if the structure was forming a synaptic specialization, then the length of the active zone(s) was delineated and measured. The active zone was defined as the length of plasma membrane directly opposing the postsynaptic density, across the synaptic cleft. Other TH-positive structures completely within the electron micrograph frame were also analyzed. Analysis of immunogold-labeled TH-positive structures was performed using PointDensity; after the perimeter of the central structure within the frame was delineated, a point marker was placed within the center of each vesicle. Vesicles were marked if at least 50% of the vesicle membrane was visible. TH-positive profiles that were forming synapses were analyzed using PointDensitySyn, the plasma membrane of the structure was traced, the active zone(s) were delineated, and points were placed in the center of the vesicles (Anwar et al., 2011; Janezic et al., 2013). Cumulative distributions for EM data were analyzed using the Kolmogorov-Smirnov test. Inter-vesicle distance data binned at 50 nm interval was analyzed using a two-way ANOVA with Sidak’s multiple comparisons test.

#### Behavior Data

Testing was performed by an observed blind to genotype. Datasets with non-normal distributions were analyzed using a Mann-Whitney test, data sets with a normal distribution were analyzed using unpaired t-tests. Rotarod and four-choice reversal test were analyzed using a two-way ANOVA with Sidak’s multiple comparisons test. The three chamber social approach test was analyzed within genotype to test for significant differences between mouse and object with paired t-tests.

## Supplemental Information

**Figure S1.**
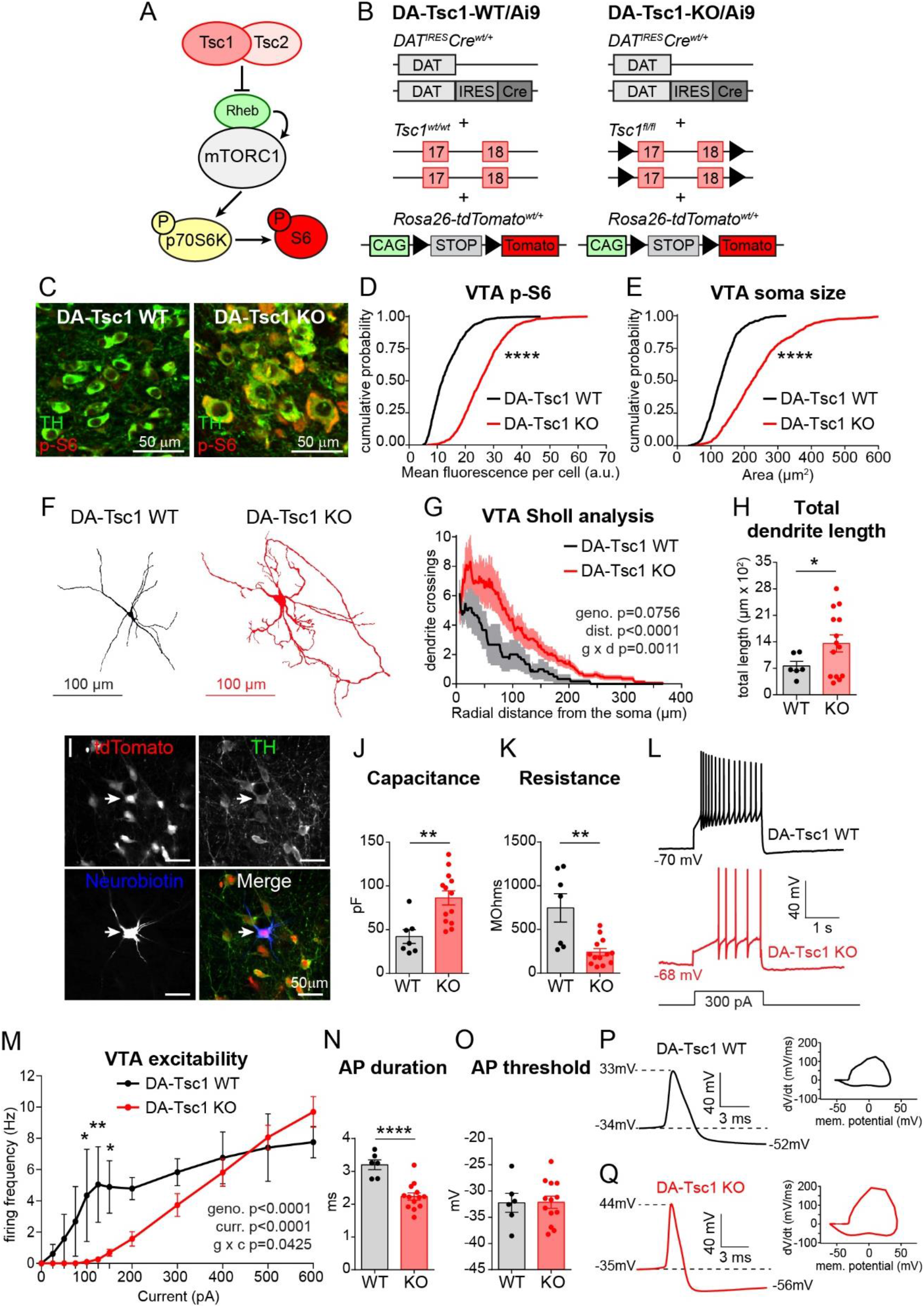
DA-Tsc1 KO VTA neurons are hypertrophic and have reduced intrinsic excitability, related to Figure 1. (A) Simplified mTORC1 signaling schematic showing the Tsc1/2 heterodimer as a negative regulator of the small GTP-ase Rheb, which directly promotes mTORC1 kinase activity. mTORC1 phosphorylates p70S6 kinase, which in turn phosphorylates ribosomal protein S6, used here as a read-out for mTORC1 activity. (B) Schematic of the genetic strategy to selectively delete *Tsc1* from DA neurons, visualized by the Ai9 tdTomato Cre-reporter. (C) Confocal images of VTA DA neurons from *DAT^IRES^Cre^wt/+^* mice homozygous for wild-type *Tsc1* (DA-Tsc1 WT, left) or floxed *Tsc1* (DA-Tsc1 KO, right). Sections were labelled with antibodies against tyrosine hydroxylase (TH) and phosphorylated S6 (p-S6, Ser240/244). (D,E) Cumulative probability plots of VTA DA neuron p-S6 levels (D) and soma area (E). Black lines show the distributions of values from DA-Tsc1 WT mice (n=892 neurons from 3 mice) and red lines show distributions for DA-Tsc1 KO mice (n=863 neurons from 3 mice). ****, p<0.0001, Kolmogorov-Smirnov test. (F) Three-dimensional reconstructions of the somata and dendrites of neurobiotin-filled VTA DA neurons from whole-cell patch-clamp experiments. (G) Sholl analysis of VTA DA neurons. Dark colored lines are the mean, the lighter color shading is SEM (DA-Tsc1 WT: n=6 neurons from 5 mice, DA-Tsc1 KO: n=14 neurons from 8 mice). Two-way ANOVA p values are shown. (H) Total dendritic length per cell measured from reconstructed VTA DA neurons. Bars represent mean ± SEM, dots represent individual neurons. (n is the same as for panel E) *, p=0.0329, unpaired, two-tailed t test. (I) Confocal images of a VTA section containing a triple-labelled (tdTomato Cre-reporter, TH, and neurobiotin) DA neuron used for three-dimensional reconstruction. (J,K) Mean ± SEM membrane capacitance (J), and membrane resistance (K). Dots indicate the values of individual neurons. (DA-Tsc1 WT: n=6 neurons from 4 mice, DA-Tsc1 KO: n=13 neurons from 8 mice). **, prersist=0.001; **, pcapacit=0.0024, unpaired, two-tailed t tests. (L) Typical examples of action potential firing elicited with a 300 pA current step in VTA DA neurons of the indicated genotypes. (M) Excitability curves showing the firing frequency of VTA DA neurons in response to two second depolarizing current steps of increasing amplitude. Data are displayed as mean ± SEM (DA-Tsc1 WT: n=7 neurons from 4 mice, DA-Tsc1 KO: n=14 neurons from 8 mice). Two-way ANOVA p values are shown. *, p100pA=0.0168; *, p150pA=0.0187; **, p125pA=0.0040; Sidak’s multiple comparisons test. (N,O) Mean ± SEM action potential (“AP”) duration (N), and membrane potential at AP threshold (O). Dots indicate the values of individual neurons. (DA-Tsc1 WT: n=6 neurons from 4 mice, DA-Tsc1 KO: n=13 neurons from 8 mice). ****, p<0.0001, unpaired, two-tailed t tests. (P,Q) Examples of individual action potentials and their respective phase plots for VTA DA neurons in DA-Tsc1 WT (P) and DA-Tsc1 KO (Q). See also Table S2 for complete electrophysiology results.

**Figure S2.**
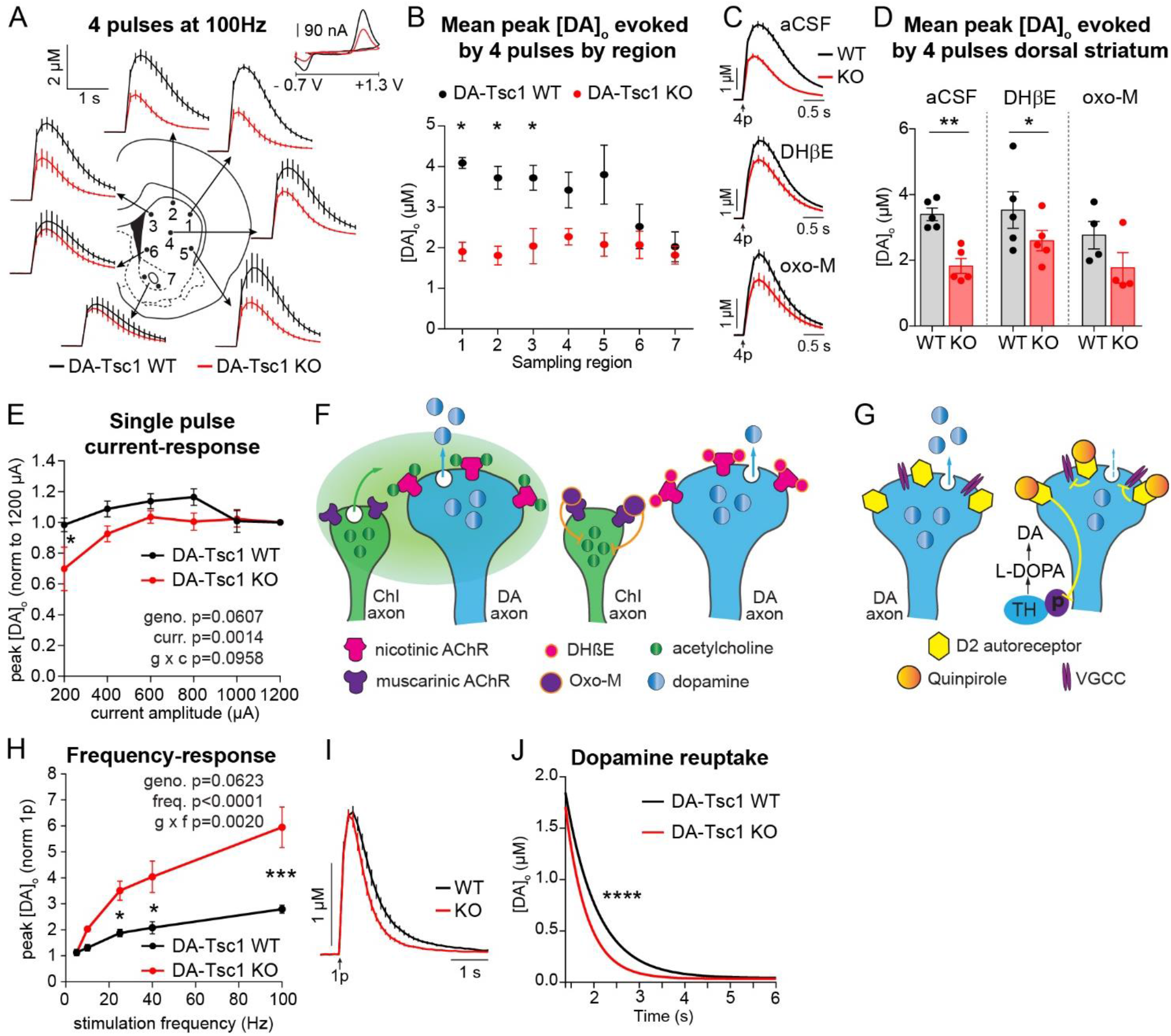
*Tsc1* deletion impairs striatal DA release and increases reuptake kinetics, related to Figure 2. (A) Mean extracellular DA release ([DA]_o_) ± SEM versus time evoked from different striatal subregions by high frequency burst stimulation (4 pulses at 100 Hz). Traces are an average of 7-10 release transients per site from 5 mice per genotype. 1-dorsolateral striatum, 2-dorsocentral striatum, 3-dorsomedial striatum, 4-central striatum, 5-ventrolateral striatum, 6-ventromedial striatum, 7-nucleus accumbens core (two sampling sites within the core were averaged together). Inset, typical cyclic voltammograms show characteristic DA waveform. (B) Mean peak high frequency stimulation-evoked [DA]_o_ ± SEM by striatal region (numbers correspond to the numbered sites in panel A). n=7-10 transients per site from 5 mice per genotype. *, p_1,2_<0.0001; *, p_3_=0.0048, paired t tests. (C) Mean high frequency stimulation-evoked [DA]_o_ ± SEM versus time for all dorsal striatum sites recorded in normal aCSF (average of 46 transients across 6 recording sites per genotype from 5 mice), DHβE (1 μM, average of 59 transients across 6 recording sites per genotype from 5 mice), or oxotremorine-M (oxo-M, 10 μM, average of 43 transients across 6 recording sites per genotype from 4 mice). (D) Mean peak high frequency stimulation-evoked [DA]_o_ ± SEM averaged across all dorsal striatum sites (sites #1-6 in panels A and B). Dots represent average evoked [DA]_o_ per mouse in aCSF, DHβE (1 μM) and oxotremorine-M (10 μM), n=4-5 mice per genotype. *, p=0.0334; **, p=0.0022, paired t tests. (E) Mean peak [DA]_o_ ± SEM evoked in the dorsomedial striatum by single pulse stimulations with current amplitude varying between 200-1200 μA delivered in pseudo-random order. Data are normalized to 1200 μA single pulse-evoked [DA]_o_ within each genotype. 600 μA stimulation intensity was used for all other experiments. n=4 mice per genotype. Two-way ANOVA p values are shown. *, p=0.0140 (for WT vs KO at 200 μA), Sidak’s multiple comparisons test. (F) Schematic showing DHßE (nicotinic AChR antagonist) and oxo-M (muscarinic AChR agonist) sites of action at nicotinic receptors on DA axon terminals and muscarinic autoreceptors on cholinergic interneurons, both of which remove cholinergic control over striatal DA transmission. (G) Schematic of D2 autoreceptor control of striatal DA release via voltage-gated calcium channels (VGCC) and tyrosine hydroxylase (TH) phosphorylation. Quinpirole is a D2 receptor agonist. (H) Mean peak [DA]_o_ ± SEM evoked by single pulse stimulation or short trains of 4 pulses at 5, 10, 25, 40 and 100 Hz delivered in pseudo-random order in dorsolateral striatum. DHβE (1 μM) applied throughout. Data are normalized to single pulse-evoked [DA]_o_ within each genotype. n=2 mice per genotype. Two-way ANOVA p values are shown. *, p25Hz=0.0254; *, p40Hz=0.0118; ***, p100Hz=0.0005, Sidak’s multiple comparisons test. (I) Mean single pulse-evoked [DA]_o_ ± SEM versus time from concentration- and region-matched FCV recordings across genotypes. Average of 10 transients from 3-4 mice per genotype. (J) Single-phase exponential decay curve-fit of the falling phase of single pulse-evoked concentration and region-matched DA transients for each genotypes. X-axis is time 375 ms after stimulation onset. n=10 traces from 3-4 mice per genotype. ****, p<0.0001, curve-fit comparison.

**Figure S3.**
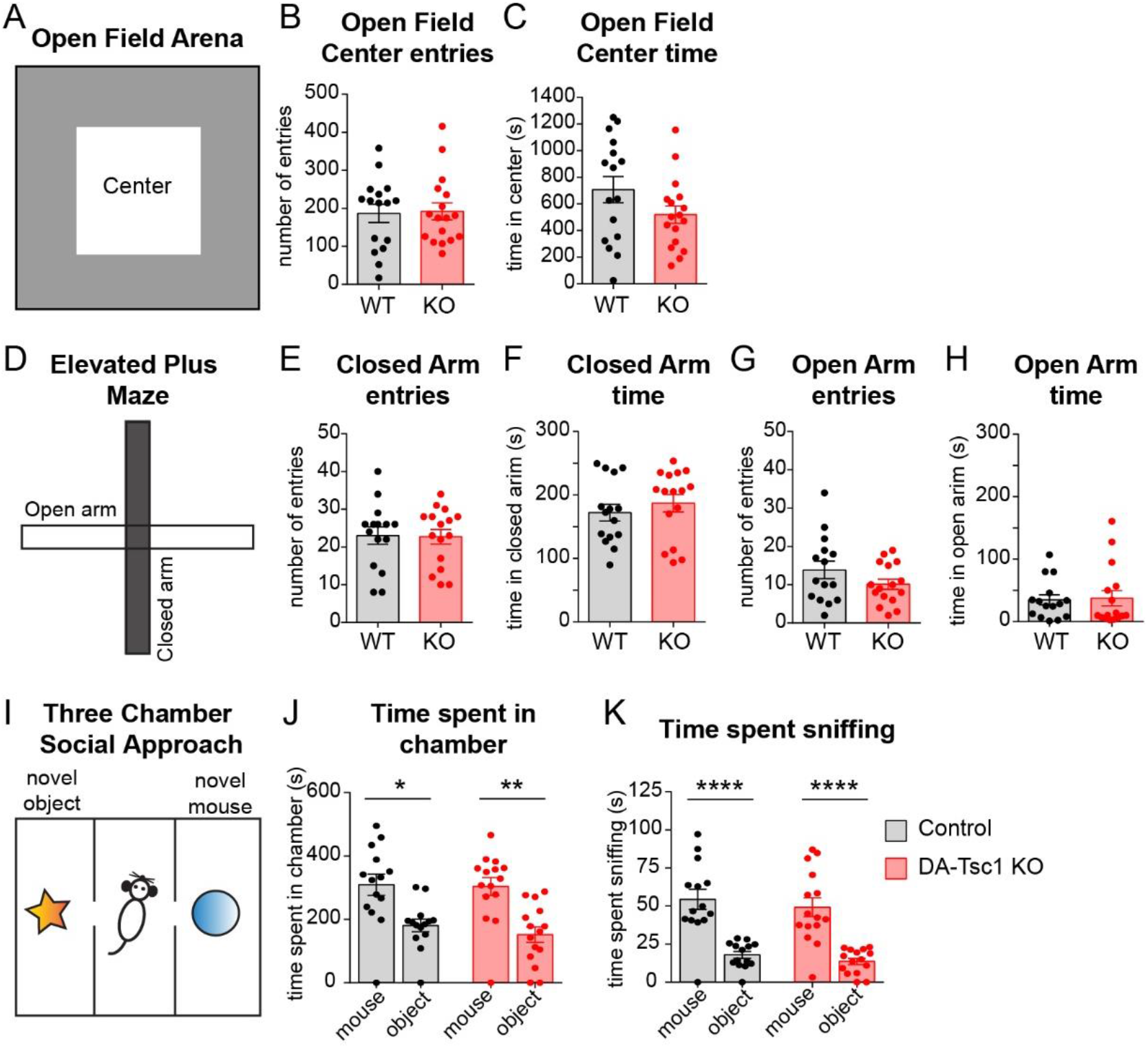
DA-Tsc1-KO mice do not show changes in anxiety or social approach, related to Figure 5. (A) Schematic of the open field arena showing the center area (white box). (B,C) Quantification of open field behavior. Mean ± SEM number of entries into the center of the arena (B) and time spent in the center (C) over 60 minutes. n=16 DA-Tsc1 WT mice and 17 DA-Tsc1 KO mice. Unpaired t tests revealed no significant differences. (D) Schematic of the elevated plus maze (EPM). (E-H) Quantification of EPM behavior. Mean ± SEM number of entries into the closed arms (E), time spent in the closed arms (F), entries into the open arms (G), and time spent in the open arms (H) over 5 minutes. n=15 DA-Tsc1 WT mice and 16 DA-Tsc1 KO mice. Unpaired t tests revealed no significant differences. (I) Schematic of the three-chamber social approach test. (J,K) Mean ± SEM time spent in the chamber with the novel mouse or novel object (J) and time spent investigating (sniffing) the novel mouse or novel object (K) during the 10 minute test. Controls are DAT^IRES^Cre negative littermates of DA-Tsc1 KO mice. Comparisons are made within genotype. n=14 control mice and 15 DA-Tsc1 KO mice. *, p=0.0192; **, p=0.0017; ****, p<0.0001; paired, two-tailed t test. For all panels, dots represent values from individual mice. See also Figure 5, Table S4 and Table S5 for behavior analysis by sex.

**Figure S4.**
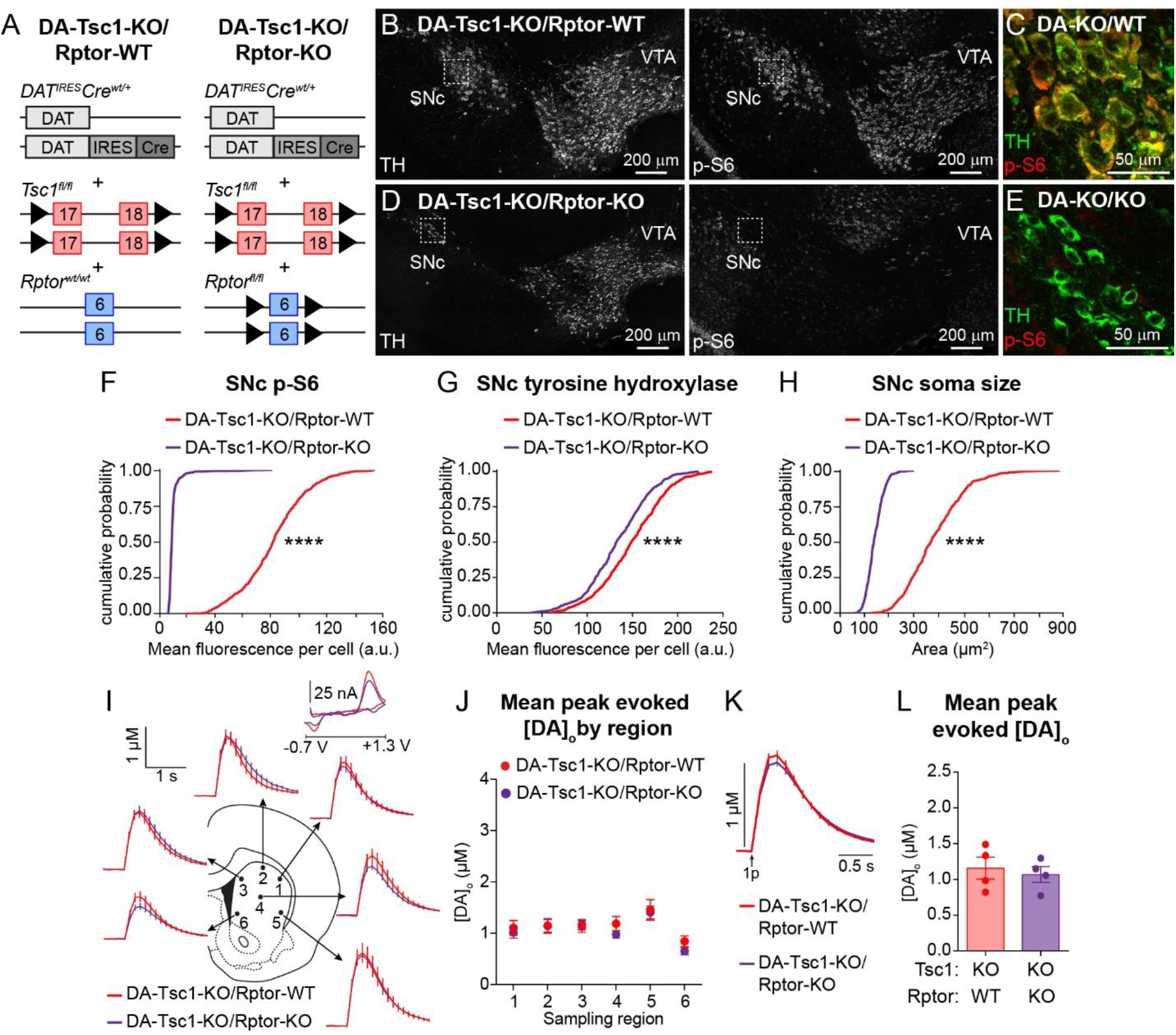
Homozygous deletion of *Rptor* from DA-Tsc1 KO neurons causes somatic hypotrophy and does not prevent dopamine release deficits, related to Figure 6. (A) Schematic of the genetic strategy to selectively delete *Tsc1* and *Rptor* in DA neurons. (B-E) Confocal images of coronal midbrain sections from DA-Tsc1-KO/Rptor-WT (B,C) and DA-Tsc1-KO/Rptor-KO (D,E) mice. Sections were labelled with antibodies against tyrosine hydroxylase (TH) and phosphorylated S6 (p-S6, Ser240/244). Panels C and E show higher magnification merged images of the boxed regions in B and D. SNc=substantia nigra pars compacta, VTA=ventral tegmental area. (F-H) Cumulative probability plots of SNc DA neuron p-S6 levels (E), tyrosine hydroxylase (TH) levels (F), and soma area (G). Red lines show the distributions of values from DA-Tsc1-KO/Rptor-WT mice (n=530 neurons from 3 mice) and green lines show distributions for DA-Tsc1-KO/Rptor-KO mice (n=669 neurons from 3 mice). ****, p<0.0001, Kolmogorov-Smirnov tests. (I) Mean extracellular DA release ([DA]_o_) ± SEM versus time evoked from different striatal subregions by single-pulse electrical stimuli. Traces show an average of 16 transients per recording site from 4 mice per genotype. 1-dorsolateral striatum, 2-dorsocentral striatum, 3-dorsomedial striatum, 4-central striatum, 5-ventrolateral striatum, 6-ventromedial striatum. Inset, typical cyclic voltammograms show characteristic DA waveform. (J) Mean peak single pulse-evoked [DA]_o_ ± SEM by striatal region (numbers correspond to the numbered sites in panel A). n=16 transients per recording site from 4 mice per genotype. Paired t tests revealed no significant difference. (K) Mean single pulse-evoked [DA]_o_ ± SEM versus time for all dorsal striatum sites recorded in normal aCSF. Average of 96 transients across 6 recording sites per genotype from 4 mice. (L) Mean peak single pulse-evoked [DA]_o_ ± SEM averaged across all dorsal striatum sites (sites #1-6 in panels A and B). Dots represent average evoked [DA]_o_ per mouse in normal aCSF. n=4 mice per genotype. Paired t test revealed no significant difference.

**Table S1.** Summary of electrophysiology data for SNc DA neurons, related to Figure 1

|  | DA-Tsc1 WT (SNc) |  |  |  | DA-Tsc1 KO (SNc) |  |  |  | Unpaired t-test (WT vs KO) |
| --- | --- | --- | --- | --- | --- | --- | --- | --- | --- |
| Properties | Mean | SEM | n (cells) | n (mice) | Mean | SEM | n (cells) | n (mice) | p-value |
| **Holding current** at -70mV (pA) | -203.06 | 28.36 | 18 | 5 | -313.23 | 20.04 | 31 | 9 | **0.0022** |
| **Series resistance** (mOhms) | 9.51 | 1.42 | 18 | 5 | 8.93 | 0.72 | 31 | 9 | 0.6875 |
| **Membrane resistance** (mOhms) | 222.83 | 30.90 | 18 | 5 | 101.97 | 8.09 | 31 | 9 | **< 0.0001** |
| **Membrane capacitance** (pF) | 64.35 | 3.82 | 18 | 5 | 128.67 | 6.28 | 31 | 9 | **< 0.0001** |
| **Resting membrane potential** (mV) | -56.13 | 1.13 | 18 | 5 | -53.62 | 2.01 | 29 | 9 | 0.3593 |
| **Current when first action potentials occur** (pA) | 145.83 | 18.66 | 18 | 5 | 341.96 | 24.70 | 28 | 9 | **< 0.0001** |
| **Action potential threshold** (mV) | -28.70 | 1.40 | 18 | 5 | -32.37 | 1.00 | 27 | 9 | **0.0338** |
| **Action potential width at threshold** (ms) | 2.77 | 0.11 | 18 | 5 | 1.94 | 0.03 | 27 | 9 | **< 0.0001** |
| **Action potential peak** (maximum membrane potential, mV) | 33.14 | 1.12 | 18 | 5 | 33.80 | 0.95 | 27 | 9 | 0.6579 |
| **Action potential height** (change in membrane potential from the start of the spike to maximum depolarization, mV) | 73.86 | 1.07 | 18 | 5 | 75.14 | 0.94 | 27 | 9 | 0.3809 |
| **Afterhyperpolarization** (minimum membrane potential after the spike, mV) | -59.08 | 1.38 | 18 | 5 | -60.89 | 0.85 | 27 | 9 | 0.2439 |
| **Afterhyperpolarization** (change in membrane potential from the start of the spike to maximum hyperpolarization, mV) | 18.36 | 0.91 | 18 | 5 | 19.56 | 0.67 | 27 | 9 | 0.2842 |

**Table S2.** Summary of electrophysiology data for VTA DA neurons, related to Figure S1

|  | DA-Tsc1 WT (VTA) |  |  |  | DA-Tsc1 KO (VTA) |  |  |  | Unpaired t-test (WT vs KO) |
| --- | --- | --- | --- | --- | --- | --- | --- | --- | --- |
| Properties | Mean | SEM | n (cells) | n (mice) | Mean | SEM | n (cells) | n (mice) | p-value |
| **Holding current** at -70mV (pA) | -117.14 | 29.34 | 7 | 4 | -189.23 | 35.40 | 13 | 8 | 0.1920 |
| **Series resistance** (mOhms) | 7.93 | 1.69 | 7 | 4 | 8.48 | 0.78 | 13 | 8 | 0.7381 |
| **Membrane resistance** (mOhms) | 746.91 | 162.31 | 7 | 4 | 241.24 | 41.31 | 13 | 8 | **0.0010** |
| **Membrane capacitance** (pF) | 42.14 | 7.90 | 7 | 4 | 86.32 | 8.10 | 13 | 8 | **0.0024** |
| **Resting membrane potential** (mV) | -55.64 | 2.19 | 7 | 4 | -57.99 | 1.40 | 12 | 8 | 0.3556 |
| **Current when first action potentials occur** (pA) | 89.29 | 23.05 | 7 | 4 | 223.08 | 31.71 | 13 | 8 | **0.0104** |
| **Action potential threshold** (mV) | -32.24 | 1.80 | 6 | 4 | -32.15 | 1.14 | 13 | 8 | 0.9659 |
| **Action potential width at threshold** (ms) | 3.20 | 0.15 | 6 | 4 | 2.23 | 0.11 | 13 | 8 | **< 0.0001** |
| **Action potential peak** (maximum membrane potential, mV) | 33.82 | 1.90 | 6 | 4 | 38.87 | 1.31 | 13 | 8 | **0.0440** |
| **Action potential height** (change in membrane potential from the start of the spike to maximum depolarization, mV) | 75.24 | 1.88 | 6 | 4 | 78.78 | 1.23 | 13 | 8 | 0.1282 |
| **Afterhyperpolarization** (minimum membrane potential after the spike, mV) | -56.99 | 2.00 | 6 | 4 | -61.32 | 1.42 | 13 | 8 | 0.1008 |
| **Afterhyperpolarization** (change in membrane potential from the start of the spike to maximum hyperpolarization, mV) | 15.57 | 1.90 | 6 | 4 | 21.42 | 0.98 | 13 | 8 | **0.0074** |

**Table S3.** Summary of electron microscopy analysis, related to Figure 4

|  | **DA-Tsc1 WT** | **DA-Tsc1 KO** | **Kolmogorov-Smirnov test (WT vs KO)** |
| --- | --- | --- | --- |
| | Mean ± SEM | Mean ± SEM | p-value |
| **All profiles** (n=239 profiles for DA-Tsc1 WT and 252 for DA-Tsc1 KO, from 4 mice per genotype) |  |  |  |
| *Perimeter ($\mu\text{m}$)* | 1.73 ± 0.05 | 2.56 ± 0.08 | **< 0.0001** |
| *Area ($\mu\text{m}^2$)* | 0.13 ± 0.005 | 0.30 ± 0.015 | **< 0.0001** |
| *Vesicles (total per profile)* | 126.3 ± 4.10 | 92.86 ± 3.73 | **< 0.0001** |
| *Vesicle density (per $\mu\text{m}^2$)* | 1217.64 ± 39.40 | 577.64 ± 35.93 | **< 0.0001** |
| *Vesicle distance to plasma membrane (nm)* | 46.04 ± 0.17 | 63.01 ± 0.31 | **< 0.0001** |
| *Inter-vesicle distance (nm)* | 273.1 ± 0.14 | 303.7 ± 0.19 | **< 0.0001** |
| *Synaptic incidence (%)* | 11.25 ± 2.50 | 10.25 ± 3.64 | > 0.9999 |
| **Profiles with post-synaptic specializations** (n=45 profiles for DA-Tsc1 WT and 41 for DA-Tsc1 KO, from 4 mice per genotype) |  |  |  |
| *Perimeter ($\mu\text{m}$)* | 1.69 ± 0.10 | 2.03 ± 0.12 | 0.1684 |
| *Area ($\mu\text{m}^2$)* | 0.12 ± 0.010 | 0.21 ± 0.024 | **0.0204** |
| *Synaptic membrane length (nm)* | 63.51 ± 3.98 | 72.31 ± 6.05 | 0.3477 |
| *Vesicles (total per profile)* | 158.7 ± 11.6 | 182.8 ± 11.32 | 0.1722 |
| *Vesicle density (per $\mu\text{m}^2$)* | 1617 ± 122.2 | 1218 ± 101.4 | 0.0716 |
| *Vesicle distance to plasma membrane (nm)* | 44.31 ± 0.35 | 57.34 ± 0.53 | **< 0.0001** |
| *Inter-vesicle distance (nm)* | 271.7 ± 0.23 | 290.3 ± 0.21 | **< 0.0001** |
| *Vesicles at the active zone (total per profile, <25nm from the post-synaptic density)* | 49.09 ± 3.73 | 42.76 ± 2.92 | 0.3509 |
| *Vesicle distance to the active zone (nm)* | 263.8 ± 0.31 | 274.3 ± 0.29 | **< 0.0001** |

**Table S4.** Summary of behavior testing results, related to Figures 5, 6, and S3

|  | **DA-Tsc1 WT** | **DA-Tsc1 KO** | **Unpaired t-test**<br>**(WT vs KO)** |
| --- | --- | --- | --- |
| | Mean ± SEM | Mean ± SEM | p-value |
| **Open Field Test** (Control n=16, KO n=17 mice) |  |  |  |
| *Distance travelled (m)* | 83.90 ± 10.05 | 99.08 ± 7.74 | 0.2415 |
| *Travelling speed (cm/s)* | 2.33 ± 0.28 | 2.76 ± 0.21 | 0.2273 |
| *Number of rears* | 366.70 ± 53.99 | 445.70 ± 48.88 | 0.2864 |
| *Time spent grooming (s)* | 134.10 ± 19.30 | 124.1 ± 21.19 | 0.7281 |
| *Time spent in the center (s)* | 706.50 ± 98.25 | 519.30 ± 64.83 | 0.1237 |
| *Number of center entries* | 186.70 ± 23.88 | 191.9 ± 22.14 | 0.8743 |
| **Elevated Plus Maze** (Control n=15, KO n=16 mice) |  |  |  |
| *Closed arm entries* | 23.00 ± 2.30 | 22.69 ± 1.95 | 0.9181 |
| *Open arm entries* | 13.87 ± 2.27 | 10.13 ± 1.32 | 0.1686 |
| *Closed arm time (s)* | 172.20 ± 13.14 | 187.10 ± 13.65 | 0.4388 |
| *Open arm time (s)* | 34.69 ± 8.12 | 37.47 ± 12.27 | 0.8519 |
|  | **Novel object** | **Novel mouse** | **Paired t-test**<br>**(within**<br>**genotype -**<br>**object vs.**<br>**mouse)** |
| | Mean ± SEM | Mean ± SEM | p-value |
| **Three Chamber Test** (Control n=14, KO n=15 mice) |  |  |  |
| *Time in chamber (s) – Control* | 180.90 ± 19.62 | 309.50 ± 33.61 | **0.0192** |
| *Time in chamber (s) – DA-Tsc1 KO* | 151.8 ± 24.30 | 303.9 ± 28.30 | **0.0017** |
| *Time spent sniffing (s) – Control* | 17.93 ± 2.24 | 54.39 ± 6.55 | **<0.0001** |
| *Time spent sniffing (s) – DA-Tsc1 KO* | 13.66 ± 2.06 | 49.18 ± 6.16 | **<0.0001** |
|  | **DA-Tsc1 WT** | **DA-Tsc1 KO** | **Sidak's**<br>**multiple**<br>**comparisons**<br>**test**<br>**(WT vs KO)** |
| | Mean ± SEM | Mean ± SEM | adjusted p-value |
| **Rotarod Test** (Control n=12, KO n=13 mice) |  |  |  |
| *Terminal speed trial 1 (rpm)* | 10.80 ± 1.25 | 13.82 ± 1.90 | 0.9989 |
| *Terminal speed trial 2 (rpm)* | 14.13 ± 1.78 | 15.80 ± 2.62 | 0.9999 |
| *Terminal speed trial 3 (rpm)* | 17.66 ± 3.23 | 16.48 ± 1.38 | 0.9999 |
| *Terminal speed trial 4 (rpm)* | 17.33 ± 2.00 | 19.25 ± 2.26 | 0.9999 |
| *Terminal speed trial 5 (rpm)* | 19.83 ± 2.33 | 20.50 ± 2.44 | 0.9999 |
| *Terminal speed trial 6 (rpm)* | 20.15 ± 2.06 | 19.64 ± 2.63 | 0.9999 |
| *Terminal speed trial 7 (rpm)* | 24.67 ± 2.63 | 23.08 ± 2.66 | 0.9999 |
| *Terminal speed trial 8 (rpm)* | 28.58 ± 2.37 | 26.54 ± 3.21 | 0.9999 |
| *Terminal speed trial 9 (rpm)* | 30.17 ± 2.04 | 28.85 ± 2.88 | 0.9999 |
| *Terminal speed trial 10 (rpm)* | 27.00 ± 2.80 | 25.00 ± 2.84 | 0.9999 |
| *Terminal speed trial 11 (rpm)* | 31.58 ± 2.63 | 27.46 ± 3.02 | 0.9640 |
| *Terminal speed trial 12 (rpm)* | 32.75 ± 3.14 | 27.00 ± 2.73 | 0.7287 |

|  | DA-Tsc1 WT | DA-Tsc1 KO | Unpaired t-test<br>(WT vs KO) |
| --- | --- | --- | --- |
|  | Mean ± SEM | Mean ± SEM | p-value |
| **Four Choice Reversal Task** (Control n=8, KO n=9 mice; males only) |  |  |  |
| *Trials to criterion - Acquisition* | 20.38 ± 2.59 | 24.78 ± 2.01 | 0.1938 |
| *Trials to criterion - Reversal* | 34.50 ± 3.19 | 45.67 ± 4.30 | 0.0594 |
| *Total errors during reversal* | 20.50 ± 2.47 | 32.22 ± 4.20 | **0.0344** |
|  | DA-Tsc1 WT | DA-Tsc1 KO | Sidak's<br>multiple<br>comparisons<br>test<br>(WT vs KO) |
|  | Mean ± SEM | Mean ± SEM | adjusted p-value |
| *Perseverative errors* | 17.25 ± 2.22 | 27.33 ± 3.67 | **0.0002** |
| *Novel errors* | 1.87 ± 0.50 | 1.00 ± 0.33 | 0.9924 |
| *Irrelevant errors* | 1.25 ± 0.41 | 1.89 ± 0.46 | 0.9977 |
| *Omissions* | 0.12 ± 0.12 | 2.00 ± 0.85 | 0.8853 |
|  | DA-Tsc1 KO/<br>Rptor WT | DA-Tsc1 KO/<br>Rptor Het | Unpaired t-test<br>(KO/WT vs<br>KO/Het) |
|  | Mean ± SEM | Mean ± SEM | p-value |
| **Four Choice Reversal Task** (KO/WT n=7, KO/HET n=10 mice; males only) |  |  |  |
| *Trials to criterion - Acquisition* | 31.00 ± 6.66 | 29.40 ± 4.33 | 0.8356 |
| *Trials to criterion - Reversal* | 49.86 ± 3.94 | 34.30 ± 4.06 | **0.0183** |
| *Total errors during reversal* | 33.43 ± 4.01 | 19.30 ± 2.55 | **0.0069** |
|  | DA-Tsc1 KO/<br>Rptor WT | DA-Tsc1 KO/<br>Rptor Het | Sidak's<br>multiple<br>comparisons<br>test<br>(KO/WT vs<br>KO/Het) |
|  | Mean ± SEM | Mean ± SEM | adjusted p-value |
| *Perseverative errors* | 27.57 ± 3.89 | 17.20 ± 2.19 | **<0.0001** |
| *Novel errors* | 1.86 ± 0.96 | 0.50 ± 0.22 | 0.9533 |
| *Irrelevant errors* | 3.29 ± 0.57 | 1.60 ± 0.62 | 0.9028 |
| *Omissions* | 0.71 ± 0.57 | 0.00 ± 0.00 | 0.9957 |

**Table S5.**
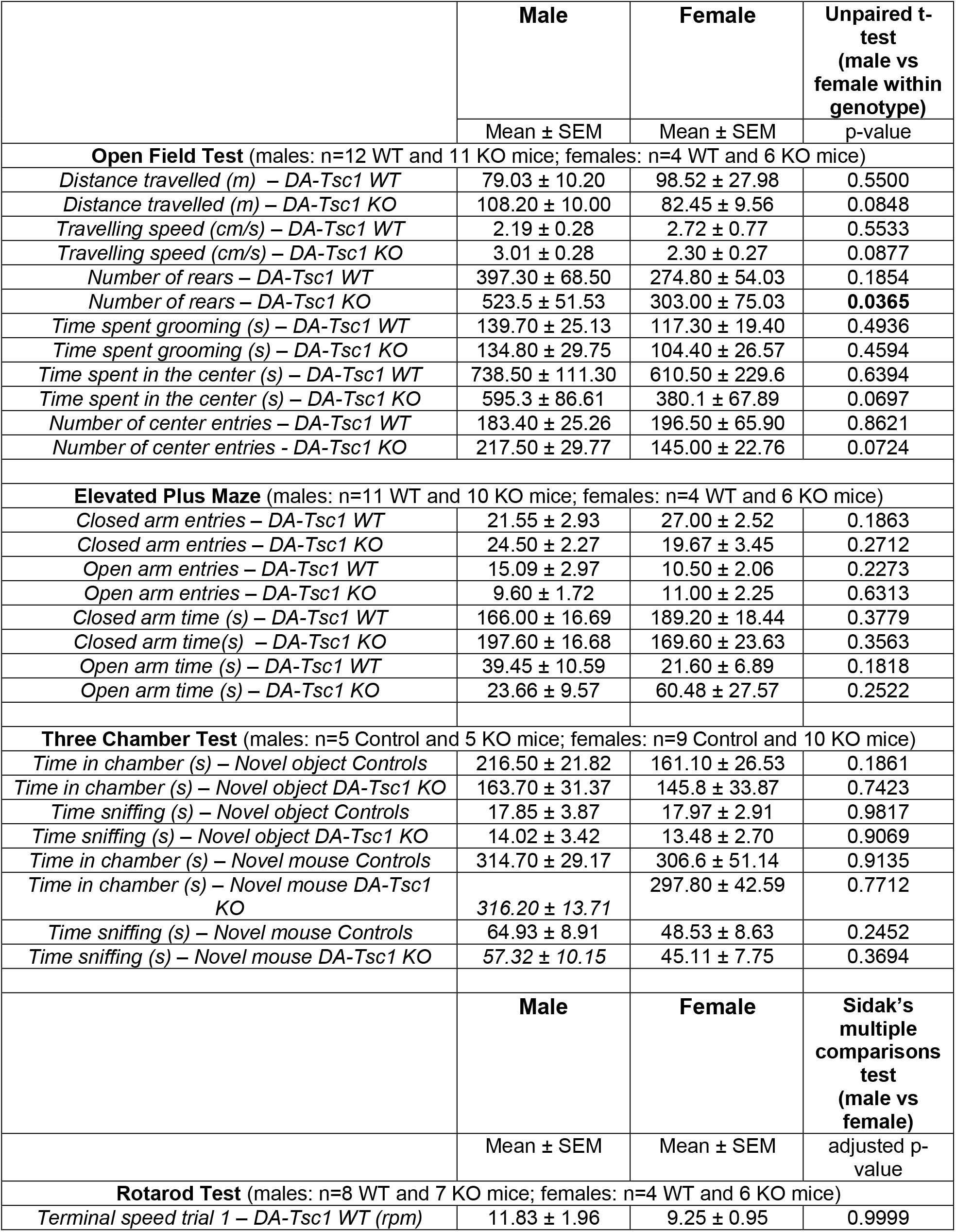
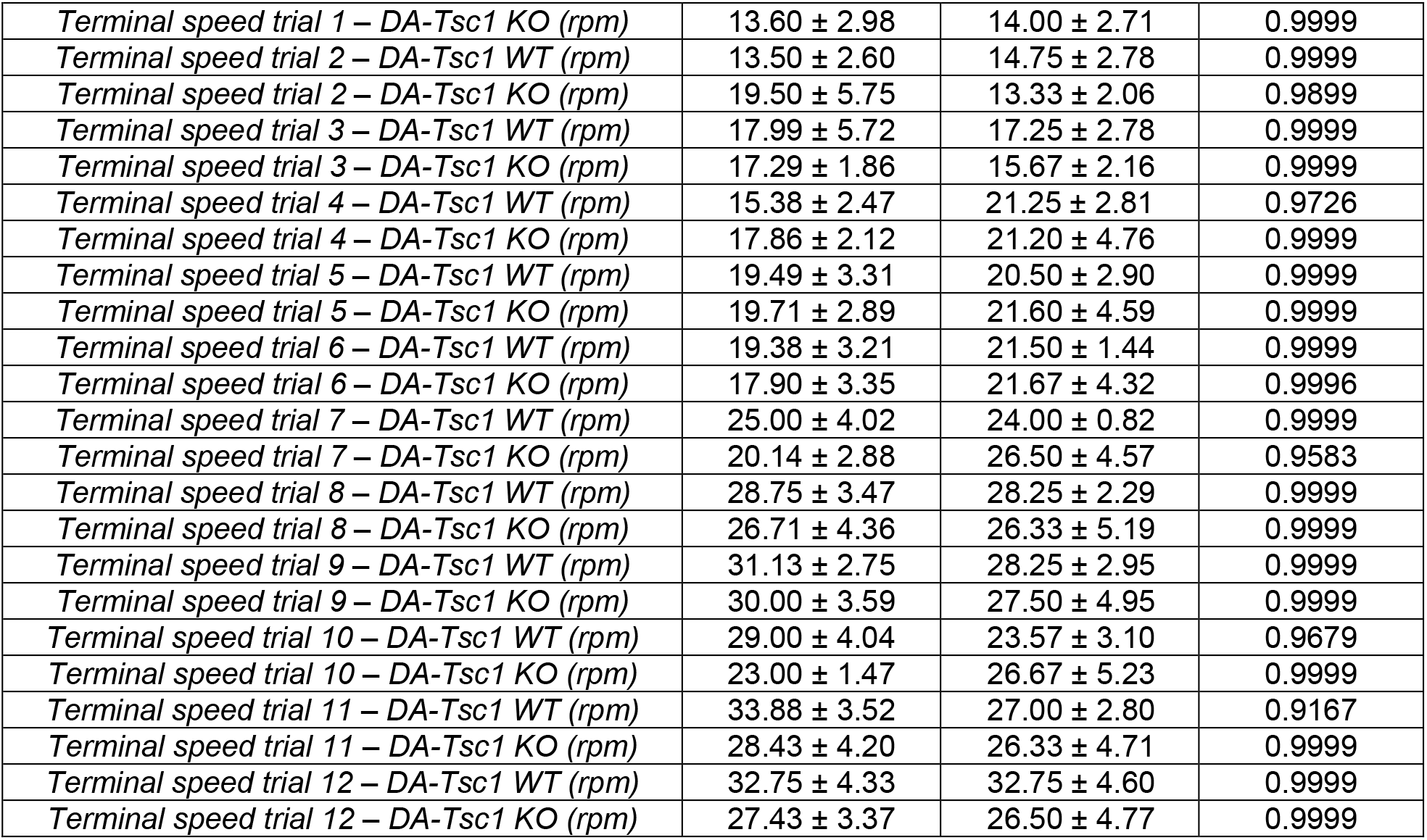
Summary of behavior testing results by sex, related to Figures 5 and S3

|  | Male | Female | Unpaired t-test<br>(male vs female within genotype) |
| --- | --- | --- | --- |
| | Mean ± SEM | Mean ± SEM | p-value |
| **Open Field Test** (males: n=12 WT and 11 KO mice; females: n=4 WT and 6 KO mice) |  |  |  |
| *Distance travelled (m) – DA-Tsc1 WT* | 79.03 ± 10.20 | 98.52 ± 27.98 | 0.5500 |
| *Distance travelled (m) – DA-Tsc1 KO* | 108.20 ± 10.00 | 82.45 ± 9.56 | 0.0848 |
| *Travelling speed (cm/s) – DA-Tsc1 WT* | 2.19 ± 0.28 | 2.72 ± 0.77 | 0.5533 |
| *Travelling speed (cm/s) – DA-Tsc1 KO* | 3.01 ± 0.28 | 2.30 ± 0.27 | 0.0877 |
| *Number of rears – DA-Tsc1 WT* | 397.30 ± 68.50 | 274.80 ± 54.03 | 0.1854 |
| *Number of rears – DA-Tsc1 KO* | 523.5 ± 51.53 | 303.00 ± 75.03 | **0.0365** |
| *Time spent grooming (s) – DA-Tsc1 WT* | 139.70 ± 25.13 | 117.30 ± 19.40 | 0.4936 |
| *Time spent grooming (s) – DA-Tsc1 KO* | 134.80 ± 29.75 | 104.40 ± 26.57 | 0.4594 |
| *Time spent in the center (s) – DA-Tsc1 WT* | 738.50 ± 111.30 | 610.50 ± 229.6 | 0.6394 |
| *Time spent in the center (s) – DA-Tsc1 KO* | 595.3 ± 86.61 | 380.1 ± 67.89 | 0.0697 |
| *Number of center entries – DA-Tsc1 WT* | 183.40 ± 25.26 | 196.50 ± 65.90 | 0.8621 |
| *Number of center entries – DA-Tsc1 KO* | 217.50 ± 29.77 | 145.00 ± 22.76 | 0.0724 |
| **Elevated Plus Maze** (males: n=11 WT and 10 KO mice; females: n=4 WT and 6 KO mice) |  |  |  |
| *Closed arm entries – DA-Tsc1 WT* | 21.55 ± 2.93 | 27.00 ± 2.52 | 0.1863 |
| *Closed arm entries – DA-Tsc1 KO* | 24.50 ± 2.27 | 19.67 ± 3.45 | 0.2712 |
| *Open arm entries – DA-Tsc1 WT* | 15.09 ± 2.97 | 10.50 ± 2.06 | 0.2273 |
| *Open arm entries – DA-Tsc1 KO* | 9.60 ± 1.72 | 11.00 ± 2.25 | 0.6313 |
| *Closed arm time (s) – DA-Tsc1 WT* | 166.00 ± 16.69 | 189.20 ± 18.44 | 0.3779 |
| *Closed arm time(s) – DA-Tsc1 KO* | 197.60 ± 16.68 | 169.60 ± 23.63 | 0.3563 |
| *Open arm time (s) – DA-Tsc1 WT* | 39.45 ± 10.59 | 21.60 ± 6.89 | 0.1818 |
| *Open arm time (s) – DA-Tsc1 KO* | 23.66 ± 9.57 | 60.48 ± 27.57 | 0.2522 |
| **Three Chamber Test** (males: n=5 Control and 5 KO mice; females: n=9 Control and 10 KO mice) |  |  |  |
| *Time in chamber (s) – Novel object Controls* | 216.50 ± 21.82 | 161.10 ± 26.53 | 0.1861 |
| *Time in chamber (s) – Novel object DA-Tsc1 KO* | 163.70 ± 31.37 | 145.8 ± 33.87 | 0.7423 |
| *Time sniffing (s) – Novel object Controls* | 17.85 ± 3.87 | 17.97 ± 2.91 | 0.9817 |
| *Time sniffing (s) – Novel object DA-Tsc1 KO* | 14.02 ± 3.42 | 13.48 ± 2.70 | 0.9069 |
| *Time in chamber (s) – Novel mouse Controls* | 314.70 ± 29.17 | 306.6 ± 51.14 | 0.9135 |
| *Time in chamber (s) – Novel mouse DA-Tsc1 KO* | 316.20 ± 13.71 | 297.80 ± 42.59 | 0.7712 |
| *Time sniffing (s) – Novel mouse Controls* | 64.93 ± 8.91 | 48.53 ± 8.63 | 0.2452 |
| *Time sniffing (s) – Novel mouse DA-Tsc1 KO* | 57.32 ± 10.15 | 45.11 ± 7.75 | 0.3694 |
|  | **Male** | **Female** | **Sidak's multiple comparisons test**<br>**(male vs female)** |
| | Mean ± SEM | Mean ± SEM | adjusted p-value |
| **Rotarod Test** (males: n=8 WT and 7 KO mice; females: n=4 WT and 6 KO mice) |  |  |  |
| *Terminal speed trial 1 – DA-Tsc1 WT (rpm)* | 11.83 ± 1.96 | 9.25 ± 0.95 | 0.9999 |
| *Terminal speed trial 1 – DA-Tsc1 KO (rpm)* | 13.60 ± 2.98 | 14.00 ± 2.71 | 0.9999 |
| *Terminal speed trial 2 – DA-Tsc1 WT (rpm)* | 13.50 ± 2.60 | 14.75 ± 2.78 | 0.9999 |
| *Terminal speed trial 2 – DA-Tsc1 KO (rpm)* | 19.50 ± 5.75 | 13.33 ± 2.06 | 0.9899 |
| *Terminal speed trial 3 – DA-Tsc1 WT (rpm)* | 17.99 ± 5.72 | 17.25 ± 2.78 | 0.9999 |
| *Terminal speed trial 3 – DA-Tsc1 KO (rpm)* | 17.29 ± 1.86 | 15.67 ± 2.16 | 0.9999 |
| *Terminal speed trial 4 – DA-Tsc1 WT (rpm)* | 15.38 ± 2.47 | 21.25 ± 2.81 | 0.9726 |
| *Terminal speed trial 4 – DA-Tsc1 KO (rpm)* | 17.86 ± 2.12 | 21.20 ± 4.76 | 0.9999 |
| *Terminal speed trial 5 – DA-Tsc1 WT (rpm)* | 19.49 ± 3.31 | 20.50 ± 2.90 | 0.9999 |
| *Terminal speed trial 5 – DA-Tsc1 KO (rpm)* | 19.71 ± 2.89 | 21.60 ± 4.59 | 0.9999 |
| *Terminal speed trial 6 – DA-Tsc1 WT (rpm)* | 19.38 ± 3.21 | 21.50 ± 1.44 | 0.9999 |
| *Terminal speed trial 6 – DA-Tsc1 KO (rpm)* | 17.90 ± 3.35 | 21.67 ± 4.32 | 0.9996 |
| *Terminal speed trial 7 – DA-Tsc1 WT (rpm)* | 25.00 ± 4.02 | 24.00 ± 0.82 | 0.9999 |
| *Terminal speed trial 7 – DA-Tsc1 KO (rpm)* | 20.14 ± 2.88 | 26.50 ± 4.57 | 0.9583 |
| *Terminal speed trial 8 – DA-Tsc1 WT (rpm)* | 28.75 ± 3.47 | 28.25 ± 2.29 | 0.9999 |
| *Terminal speed trial 8 – DA-Tsc1 KO (rpm)* | 26.71 ± 4.36 | 26.33 ± 5.19 | 0.9999 |
| *Terminal speed trial 9 – DA-Tsc1 WT (rpm)* | 31.13 ± 2.75 | 28.25 ± 2.95 | 0.9999 |
| *Terminal speed trial 9 – DA-Tsc1 KO (rpm)* | 30.00 ± 3.59 | 27.50 ± 4.95 | 0.9999 |
| *Terminal speed trial 10 – DA-Tsc1 WT (rpm)* | 29.00 ± 4.04 | 23.57 ± 3.10 | 0.9679 |
| *Terminal speed trial 10 – DA-Tsc1 KO (rpm)* | 23.00 ± 1.47 | 26.67 ± 5.23 | 0.9999 |
| *Terminal speed trial 11 – DA-Tsc1 WT (rpm)* | 33.88 ± 3.52 | 27.00 ± 2.80 | 0.9167 |
| *Terminal speed trial 11 – DA-Tsc1 KO (rpm)* | 28.43 ± 4.20 | 26.33 ± 4.71 | 0.9999 |
| *Terminal speed trial 12 – DA-Tsc1 WT (rpm)* | 32.75 ± 4.33 | 32.75 ± 4.60 | 0.9999 |
| *Terminal speed trial 12 – DA-Tsc1 KO (rpm)* | 27.43 ± 3.37 | 26.50 ± 4.77 | 0.9999 |

## References

Anwar, S., Peters, O., Millership, S., Ninkina, N., Doig, N., Connor-Robson, N., Threlfell, S., Kooner, G., Deacon, R.M., Bannerman, D.M., et al. (2011). Functional alterations to the nigrostriatal system in mice lacking all three members of the synuclein family. J. Neurosci. 31, 7264–7274.

Bäckman, C.M., Malik, N., Zhang, Y., Shan, L., Grinberg, A., Hoffer, B.J., Westphal, H., and Tomac, A.C. (2006). Characterization of a mouse strain expressing Cre recombinase from the 3’ untranslated region of the dopamine transporter locus. Genesis 44, 383–390.

Bateup, H.S., Takasaki, K.T., Saulnier, J.L., Denefrio, C.L., and Sabatini, B.L. (2011). Loss of Tsc1 in vivo impairs hippocampal mGluR-LTD and increases excitatory synaptic function. J. Neurosci. 31, 8862–8869.

Bateup, H.S., Johnson, C.A., Denefrio, C.L., Saulnier, J.L., Kornacker, K., and Sabatini, B.L. (2013). Excitatory/inhibitory synaptic imbalance leads to hippocampal hyperexcitability in mouse models of tuberous sclerosis. Neuron 78, 510–522.

Bean, B.P. (2007). The action potential in mammalian central neurons. Nat. Rev. Neurosci. 8, 451–465.

Benthall, K.N., Ong, S.L., and Bateup, H.S. (2018). Corticostriatal Transmission Is Selectively Enhanced in Striatonigral Neurons with Postnatal Loss of Tsc1. Cell Rep. 23, 3197–3208.

Berke, J.D. (2018). What does dopamine mean? Nat. Neurosci. 21, 787–793.

Cartier, E.A., Parra, L.A., Baust, T.B., Quiroz, M., Salazar, G., Faundez, V., Egaña, L., and Torres, G.E. (2010). A biochemical and functional protein complex involving dopamine synthesis and transport into synaptic vesicles. J. Biol. Chem. 285, 1957–1966.

Cheng, H.-C., Kim, S.R., Oo, T.F., Kareva, T., Yarygina, O., Rzhetskaya, M., Wang, C., During, M., Talloczy, Z., Tanaka, K., et al. (2011). Akt suppresses retrograde degeneration of dopaminergic axons by inhibition of macroautophagy. J. Neurosci. 31, 2125–2135.

Cheng, H.C., Ulane, C.M., and Burke, R.E. (2010). Clinical progression in Parkinson disease and the neurobiology of axons. Ann. Neurol. 67, 715–725.

Collo, G., Bono, F., Cavalleri, L., Plebani, L., Merlo Pich, E., Millan, M.J., Spano, P.F., and Missale, C. (2012). Pre-synaptic dopamine D3 receptor mediates cocaine-induced structural plasticity in mesencephalic dopaminergic neurons via ERK and Akt pathways. J. Neurochem. 120, 765–778.

Collo, G., Bono, F., Cavalleri, L., Plebani, L., Mitola, S., Merlo Pich, E., Millan, M.J., Zoli, M., Maskos, U., Spano, P., et al. (2013). Nicotine-induced structural plasticity in mesencephalic dopaminergic neurons is mediated by dopamine D3 receptors and Akt-mTORC1 signaling. Mol. Pharmacol. 83, 1176–1189.

Cools, R., Barker, R.A., Sahikian, B.J., and Robbins, T.W. (2001). Mechanisms of cognitive set flexibility in Parkinson’s disease. Brain 124, 2503–2512.

Costa-Mattioli, M., and Monteggia, L.M. (2013). mTOR complexes in neurodevelopmental and neuropsychiatric disorders. Nat. Neurosci. 16, 1537–1543.

Cragg, S.J. (2003). Variable Dopamine Release Probability and Short-Term Plasticity Between Functional Domains of the Primate Striatum. Neuroscience 23, 4378–4385.

Cragg, S.J., and Rice, M.E. (2004). DAncing past the DAT at a DA synapse. Trends Neurosci. 27, 270–277.

D’Cruz, A.-M., Ragozzino, M.E., Mosconi, M.W., Shrestha, S., Cook, E.H., and Sweeney, J.A. (2013). Reduced behavioral flexibility in autism spectrum disorders. Neuropsychology 27, 152–160.

Dajani, D.R., and Uddin, L.Q. (2015). Demystifying cognitive flexibility: Implications for clinical and developmental neuroscience. Trends Neurosci. 38, 571–578.

Darvas, M., and Palmiter, R.D. (2011). Contributions of striatal dopamine signaling to the modulation of cognitive flexibility. Biol. Psychiatry 69, 704–707.

Dayas, C. V., Smith, D.W., and Dunkley, P.R. (2012). An emerging role for the mammalian target of rapamycin in “pathological” protein translation: relevance to cocaine addiction. Front. Pharmacol. 3, 1–12.

Diaz-Ruiz, O., Zapata, A., Shan, L., Zhang, Y., Tomac, A.C., Malik, N., de la Cruz, F., and Bäckman, C.M. (2009). Selective deletion of PTEN in dopamine neurons leads to trophic effects and adaptation of striatal medium spiny projecting neurons. PLoS One 4, 1–13.

Doig, N.M., Moss, J., and Bolam, J.P. (2010). Cortical and thalamic innervation of direct and indirect pathway medium-sized spiny neurons in mouse striatum. J. Neurosci. 30, 14610–14618.

Doig, N.M., Magill, P.J., Apicella, P., Bolam, J.P., and Sharott, A. (2014). Cortical and Thalamic Excitation Mediate the Multiphasic Responses of Striatal Cholinergic Interneurons to Motivationally Salient Stimuli. J. Neurosci. 34, 3101–3117.

Domanskyi, A., Geissler, C., Vinnikov, I.A., Alter, H., Schober, A., Vogt, M.A., Gass, P., Parlato, R., and Schutz, G. (2011). Pten ablation in adult dopaminergic neurons is neuroprotective in Parkinson’s disease models. FASEB J. 25, 2898–2910.

Ehninger, D., Han, S., Shilyansky, C., Zhou, Y., Li, W., Kwiatkowski, D.J., Ramesh, V., and Silva, A.J. (2008). Reversal of learning deficits in a Tsc2+/-mouse model of tuberous sclerosis. Nat. Med. 14, 843–848.

Gantz, S.C., Ford, C.P., Morikawa, H., and Williams, J.T. (2018). The Evolving Understanding of Dopamine Neurons in the Substantia Nigra and Ventral Tegmental Area. Annu. Rev. Physiol. 80, 219–241.

Goto, J., Talos, D.M., Klein, P., Qin, W., Chekaluk, Y.I., Anderl, S., Malinowska, I.A., Di Nardo, A., Bronson, R.T., Chan, J.A., et al. (2011). Regulable neural progenitor-specific Tsc1 loss yields giant cells with organellar dysfunction in a model of tuberous sclerosis complex. Proc. Natl. Acad. Sci. 108, E1070–E1079.

Grospe, G.M., Baker, P.M., and Ragozzino, M.E. (2018). Cognitive Flexibility Deficits Following 6-OHDA Lesions of the Rat Dorsomedial Striatum. Neuroscience 374, 80–90.

Hernandez, D., Torres, C.A., Setlik, W., Cebrián, C., Mosharov, E. V., Tang, G., Cheng, H.C., Kholodilov, N., Yarygina, O., Burke, R.E., et al. (2012). Regulation of Presynaptic Neurotransmission by Macroautophagy. Neuron 74, 277–284.

Hoeffer, C.A., and Klann, E. (2010). mTOR signaling: At the crossroads of plasticity, memory and disease. Trends Neurosci. 33, 67–75.

Hoffman, A.F., Lupica, C.R., and Gerhardt, G.A. (1998). Dopamine transporter activity in the substantia nigra and striatum assessed by high-speed chronoamperometric recordings in brain slices. J. Pharmacol. Exp. Ther. 287, 487–496.

Huang, J., and Manning, B.D. (2008). The TSC1–TSC2 complex: a molecular switchboard controlling cell growth. Biochem. J. 412, 179–190.

Janezic, S., Threlfell, S., Dodson, P.D., Dowie, M.J., Taylor, T.N., Potgieter, D., Parkkinen, L., Senior, S.L., Anwar, S., Ryan, B., et al. (2013). Deficits in dopaminergic transmission precede neuron loss and dysfunction in a new Parkinson model. Proc. Natl. Acad. Sci. 110, E4016–E4025.

Johnson, C.M., Peckler, H., Tai, L.H., and Wilbrecht, L. (2016). Rule learning enhances structural plasticity of long-range axons in frontal cortex. Nat. Commun. 7, 1–14.

Kim, D.-H., Sarbassov, D.D., Ali, S.M., King, J.E., Latek, R.R., Erdjument-Bromage, H., Tempst, P., and Sabatini, D.M. (2002). mTOR interacts with raptor to form a nutrient-sensitive complex that signals to the cell growth machinery. Cell 110, 163–175.

Kim, S.R., Kareva, T., Yarygina, O., Kholodilov, N., and Burke, R.E. (2012). AAV transduction of dopamine neurons with constitutively active rheb protects from neurodegeneration and mediates axon regrowth. Mol. Ther. 20, 275–286.

Kimm, T., Khaliq, Z.M., and Bean, B.P. (2015). Differential Regulation of Action Potential Shape and Burst-Frequency Firing by BK and Kv2 Channels in Substantia Nigra Dopaminergic Neurons. J. Neurosci. 35, 16404–16417.

Klanker, M., Feenstra, M., and Denys, D. (2013). Dopaminergic control of cognitive flexibility in humans and animals. Front. Neurosci. 7, 1–24.

Kwiatkowski, D.J., Zhang, H., Bandura, J.L., Heiberger, K.M., Glogauer, M., el-Hashemite, N., and Onda, H. (2002). A mouse model of TSC1 reveals sex-dependent lethality from liver hemangiomas, and up-regulation of p70S6 kinase activity in Tsc1 null cells. Hum. Mol. Genet. 11, 525–534.

Kwon, C.H., Luikart, B.W., Powell, C.M., Zhou, J., Matheny, S.A., Zhang, W., Li, Y., Baker, S.J., and Parada, L.F. (2006). Pten Regulates Neuronal Arborization and Social Interaction in Mice. Neuron 50, 377–388.

Lan, A., Chen, J., Zhao, Y., Chai, Z., and Hu, Y. (2017). mTOR Signaling in Parkinson’s Disease. NeuroMolecular Med. 19, 1–10.

Laplante, M., and Sabatini, D.M. (2012). mTOR signaling in growth control and disease. Cell 149, 274–293.

Lipton, J.O., and Sahin, M. (2014). The Neurology of mTOR. Neuron 84, 275–291.

Liu, C., Kershberg, L., Wang, J., Schneeberger, S., and Kaeser, P.S. (2018a). Dopamine Secretion Is Mediated by Sparse Active Zone-like Release Sites. Cell 172, 1–13.

Liu, X., Li, Y., Yu, L., Vickstrom, C.R., and Liu, Q. (2018b). VTA mTOR Signaling Regulates Dopamine Dynamics, Cocaine-Induced Synaptic Alterations and Reward. Neuropsychopharmacology 43, 1066–1077.

Luo, S.X., Timbang, L., Kim, J.-I., Shang, Y., Sandoval, K., Tang, A.A., Whistler, J.L., Ding, J.B., and Huang, E.J. (2016). TGF-β Signaling in Dopaminergic Neurons Regulates Dendritic Growth, Excitatory-Inhibitory Synaptic Balance, and Reversal Learning. Cell Rep. 17, 3233–3245.

Madisen, L., Zwingman, T.A., Sunkin, S.M., Oh, S.W., Zariwala, H.A., Gu, H., Ng, L.L., Palmiter, R.D., Hawrylycz, M.J., Jones, A.R., et al. (2010). A robust and high-throughput Cre reporting and characterization system for the whole mouse brain. Nat. Neurosci. 13, 133–140.

Malagelada, C., Jin, Z.H., Jackson-Lewis, V., Przedborski, S., and Greene, L.A. (2010). Rapamycin protects against neuron death in in vitro and in vivo models of Parkinson’s disease. J. Neurosci. 30, 1166–1175.

Manduca, A., Servadio, M., Damsteegt, R., Campolongo, P., Vanderschuren, L.J.M.J., and Trezza, V. (2016). Dopaminergic neurotransmission in the nucleus accumbens modulates social play behavior in rats. Neuropsychopharmacology 41, 2215–2223.

Matsuda, W., Furuta, T., Nakamura, K.C., Hioki, H., Fujiyama, F., Arai, R., and Kaneko, T. (2009). Single nigrostriatal dopaminergic neurons form widely spread and highly dense axonal arborizations in the neostriatum. J. Neurosci. 29, 444–453.

Mazei-Robison, M.S., Koo, J.W., Friedman, A.K., Lansink, C.S., Robison, A.J., Vinish, M., Krishnan, V., Kim, S., Siuta, M.A., Galli, A., et al. (2011). Role for mTOR signaling and neuronal activity in morphine-induced adaptations in ventral tegmental area dopamine neurons. Neuron 72, 977–990.

Montague, P.R., McClure, S.M., Baldwin, P.R., Phillips, P.E.M., Budygin, E.A., Stuber, G.D., Kilpatrick, M.R., and Wightman, R.M. (2004). Dynamic gain control of dopamine delivery in freely moving animals. J. Neurosci. 24, 1754–1759.

Moss, J., and Bolam, J.P. (2008). A dopaminergic axon lattice in the striatum and its relationship with cortical and thalamic terminals. J. Neurosci. 28, 11221–11230.

Neasta, J., Barak, S., Hamida, S. Ben, and Ron, D. (2014). MTOR complex 1: A key player in neuroadaptations induced by drugs of abuse. J. Neurochem. 130, 172–184.

Nirenberg, M.J., Vaughan, R.A., Uhl, G.R., Kuhar, M.J., and Pickel, V.M. (1996). The dopamine transporter is localized to dendritic and axonal plasma membranes of nigrostriatal dopaminergic neurons. J. Neurosci. 16, 436–447.

Normand, E.A., Crandall, S.R., Thorn, C.A., Murphy, E.M., Voelcker, B., Browning, C., Machan, J., Moore, C.I., Connors, B.W., and Zervas, M. (2013). Temporal and mosaic Tsc1 Deletion in the developing thalamus disrupts thalamocortical circuitry, neural function, and behavior. Neuron 78, 895–909.

O’Neill, M., and Brown, V.J. (2007). The effect of striatal dopamine depletion and the adenosine A2A antagonist KW-6002 on reversal learning in rats. Neurobiol. Learn. Mem. 88, 75–81.

Park, H., Li, Y., and Tsien, R.W. (2012). Influence of Synaptic Vesicle Position on Release Probability and Exocytotic Fusion Mode. Science 335, 1362–1366.

Paxinos, G., and Franklin, K.B.J. (2008). The Mouse Brain in Stereotaxic Coordinates (New York: Elsevier Academic Press).

Potter, W.B., Basu, T., O’Riordan, K.J., Kirchner, A., Rutecki, P., Burger, C., and Roopra, A. (2013). Reduced Juvenile Long-Term Depression in Tuberous Sclerosis Complex Is Mitigated in Adults by Compensatory Recruitment of mGluR5 and Erk Signaling. PLoS Biol. 11, 1–12.

Prather, P., and de Vries, P.J. (2004). Behavioral and cognitive aspects of tuberous sclerosis complex. J. Child Neurol. 19, 666–674.

Raab-Graham, K.F., Haddick, P.C.G., Jan, Y.N., and Jan, L.Y. (2006). mRNA Translation in Dendrites. Science. 314, 144–148.

Rice, M.E., and Cragg, S.J. (2004). Nicotine amplifies reward-related dopamine signals in striatum. Nat. Neurosci. 7, 583–584.

Rice, M.E., and Cragg, S.J. (2008). Dopamine spillover after quantal release: rethinking dopamine transmission in the nigrostriatal pathway. Brain Res. Rev. 58, 303–313.

Rice, M.E., Patel, J.C., and Cragg, S.J. (2011). Dopamine release in the basal ganglia. Neuroscience 198, 112–137.

Rothwell, P.E., Fuccillo, M. V, Maxeiner, S., Hayton, S.J., Gokce, O., Lim, B.K., Fowler, S.C., Malenka, R.C., and Südhof, T.C. (2014). Autism-associated neuroligin-3 mutations commonly impair striatal circuits to boost repetitive behaviors. Cell 158, 198–212.

Santini, E., Huynh, T.N., MacAskill, A.F., Carter, A.G., Pierre, P., Ruggero, D., Kaphzan, H., and Klann, E. (2013). Exaggerated translation causes synaptic and behavioural aberrations associated with autism. Nature 493, 411–415.

Schultz, W. (2005). Behavioral Theories and the Neurophysiology of Reward. Annu. Rev. Psychol. 57, 87–115.

Sengupta, S., Peterson, T.R., Laplante, M., Oh, S., and Sabatini, D.M. (2010). MTORC1 controls fasting-induced ketogenesis and its modulation by ageing. Nature 468, 1100–1106.

Sergeant, J.A., Geurts, H., and Oosterlaan, J. (2002). How specific is a deficit of executive functioning for attention-deficit/ hyperactivity disorder. Behav. Brain Res. 130, 3–28.

Steinberg, E.E., Keiflin, R., Boivin, J.R., Witten, I.B., Deisseroth, K., and Janak, P.H. (2013). A causal link between prediction errors, dopamine neurons and learning. Nat. Neurosci. 16, 966–973.

Sulzer, D., Cragg, S.J., and Rice, M.E. (2016). Striatal dopamine neurotransmission: regulation of release and uptake. Basal Ganglia 6, 123–148.

Tavazoie, S.F., Alvarez, V.A., Ridenour, D.A., Kwiatkowski, D.J., and Sabatini, B.L. (2005). Regulation of neuronal morphology and function by the tumor suppressors Tsc1 and Tsc2. Nat. Neurosci. 8, 1727–1734.

Threlfell, S., Clements, M.A., Khodai, T., Pienaar, I.S., Exley, R., Wess, J., and Cragg, S.J. (2010). Striatal muscarinic receptors promote activity dependence of dopamine transmission via distinct receptor subtypes on cholinergic interneurons in ventral versus dorsal striatum. J. Neurosci. 30, 3398–3408.

Threlfell, S., Lalic, T., Platt, N.J., Jennings, K.A., Deisseroth, K., and Cragg, S.J. (2012). Striatal dopamine release is triggered by synchronized activity in cholinergic interneurons. Neuron 75, 58–64.

Tsai, P.T., Hull, C., Chu, Y., Greene-Colozzi, E., Sadowski, A.R., Leech, J.M., Steinberg, J., Crawley, J.N., Regehr, W.G., and Sahin, M. (2012). Autistic-like behaviour and cerebellar dysfunction in Purkinje cell Tsc1 mutant mice. Nature 488, 647–651.

de Vries, P.J., Whittemore, V.H., Leclezio, L., Byars, A.W., Dunn, D., Ess, K.C., Hook, D., King, B. H., Sahin, M., and Jansen, A. (2015). Tuberous sclerosis associated neuropsychiatric disorders (TAND) and the TAND Checklist. Pediatr. Neurol. 52, 25–35.

Walf, A.A., and Frye, C.A. (2007). The use of the elevated plus maze as an assay of anxiety-related behavior in rodents. Nat. Protoc. 2, 322–328.

Watson, G., and Leverenz, J.B. (2010). Profile of Cognitive Impairment in Parkinson’s Disease. Brain Pathol. 20, 640–645.

Weston, M.C., Chen, H., and Swann, J.W. (2014). Loss of mTOR repressors Tsc1 or Pten has divergent effects on excitatory and inhibitory synaptic transmission in single hippocampal neuron cultures. Front. Mol. Neurosci. 7, 1–15.

Wickens, J.R., Horvitz, J.C., Costa, R.M., and Killcross, S. (2007). Dopaminergic mechanisms in actions and habits. J. Neurosci. 27, 8181–8183.

Williams-Gray, C.H., Foltynie, T., Brayne, C.E.G., Robbins, T.W., and Barker, R.A. (2007). Evolution of cognitive dysfunction in an incident Parkinson’s disease cohort. Brain 130, 1787–1798.

Wu, J., McCallum, S.E., Glick, S.D., and Huang, Y. (2011). Inhibition of the mammalian target of rapamycin pathway by rapamycin blocks cocaine-induced locomotor sensitization. Neuroscience 172, 104–109.

Xu, Y., Liu, C., Chen, S., Ye, Y., Guo, M., Ren, Q., Liu, L., Zhang, H., Xu, C., Zhou, Q., et al. (2014). Activation of AMPK and inactivation of Akt result in suppression of mTOR-mediated S6K1 and 4E-BP1 pathways leading to neuronal cell death in in vitro models of Parkinson’s disease. Cell. Signal. 26, 1680–1689.

Yang, M., Silverman, J.L., and Crawley, J.N. (2011). Automated three-chambered social approach task for mice. Curr. Protoc. Neurosci. 56, 1–23.

Yang, S.-B., Tien, A.-C., Boddupalli, G., Xu, A.W., Jan, Y.N., and Jan, L.Y. (2012). Rapamycin ameliorates age-dependent obesity associated with increased mTOR signaling in hypothalamic POMC neurons. Neuron 75, 425–436.

Zhang, H., and Sulzer, D. (2012). Basal Ganglia. 2, 5–13.

Zhou, Q., Liu, C., Liu, W., Zhang, H., Zhang, R., Liu, J., Zhang, J., Xu, C., Liu, L., Huang, S., et al. (2015). Rotenone induction of hydrogen peroxide inhibits mTOR-mediated S6K1 and 4E-BP1/eIF4E pathways, leading to neuronal apoptosis. Toxicol. Sci. 143, 81–96.

Zweifel, L.S., Fadok, J.P., Argilli, E., Garelick, M.G., Jones, G.L., Dickerson, T.M.K., Allen, J.M., Mizumori, S.J.Y., Bonci, A., and Palmiter, R.D. (2011). Activation of dopamine neurons is critical for aversive conditioning and prevention of generalized anxiety. Nat. Neurosci. 14, 620–626.

